# Cells transit through a quiescent-like state to convert to neurons at high rates

**DOI:** 10.1101/2024.11.22.624928

**Authors:** A Beitz, JMY Teves, C Oakes, C Johnstone, N Wang, JM Brickman, KE Galloway

## Abstract

While transcription factors (TFs) provide essential cues for directing and redirecting cell fate, TFs alone are insufficient to drive cells to adopt alternative fates. Rather, transcription factors rely on receptive cell states to induce novel identities. Cell state emerges from and is shaped by cellular history and the activity of diverse processes. Here, we define the cellular and molecular properties of a highly receptive state amenable to transcription factor-mediated direct conversion from fibroblasts to induced motor neurons. Using a well-defined model of direct conversion to a post-mitotic fate, we identify the highly proliferative, receptive state that transiently emerges during conversion. Through examining chromatin accessibility, histone marks, and nuclear features, we find that cells reprogram from a state characterized by global reductions in nuclear size and transcriptional activity. Supported by globally increased levels of H3K27me3, cells enter a quiescent-like state of reduced RNA metabolism and elevated expression of REST and p27, markers of quiescent neural stem cells. From this transient state, cells convert to neurons at high rates. Inhibition of Ezh2, the catalytic subunit of PRC2 that deposits H3K27me3, abolishes conversion. Our work offers a roadmap to identify global changes in cellular processes that define cells with different conversion potentials that may generalize to other cell-fate transitions.

**Highlights:**

- Proliferation drives cells to a compact nuclear state that is receptive to TF-mediated conversion.
- Increased receptivity to TFs corresponds to reduced nuclear volumes.
- Reprogrammable cells display global, genome-wide increases in H3K27me3.
- High levels of H3K27me3 support cells’ transits through a state of altered RNA metabolism.
- Inhibition of Ezh2 increases nuclear size, reduces the expression of the quiescence marker p27.
- Acute inhibition of Ezh2 abolishes motor neuron conversion.

**One Sentence Summary:** Cells transit through a quiescent-like state characterized by global reductions in nuclear size and transcriptional activity to convert to neurons at high rates.

**Graphical Abstract:** 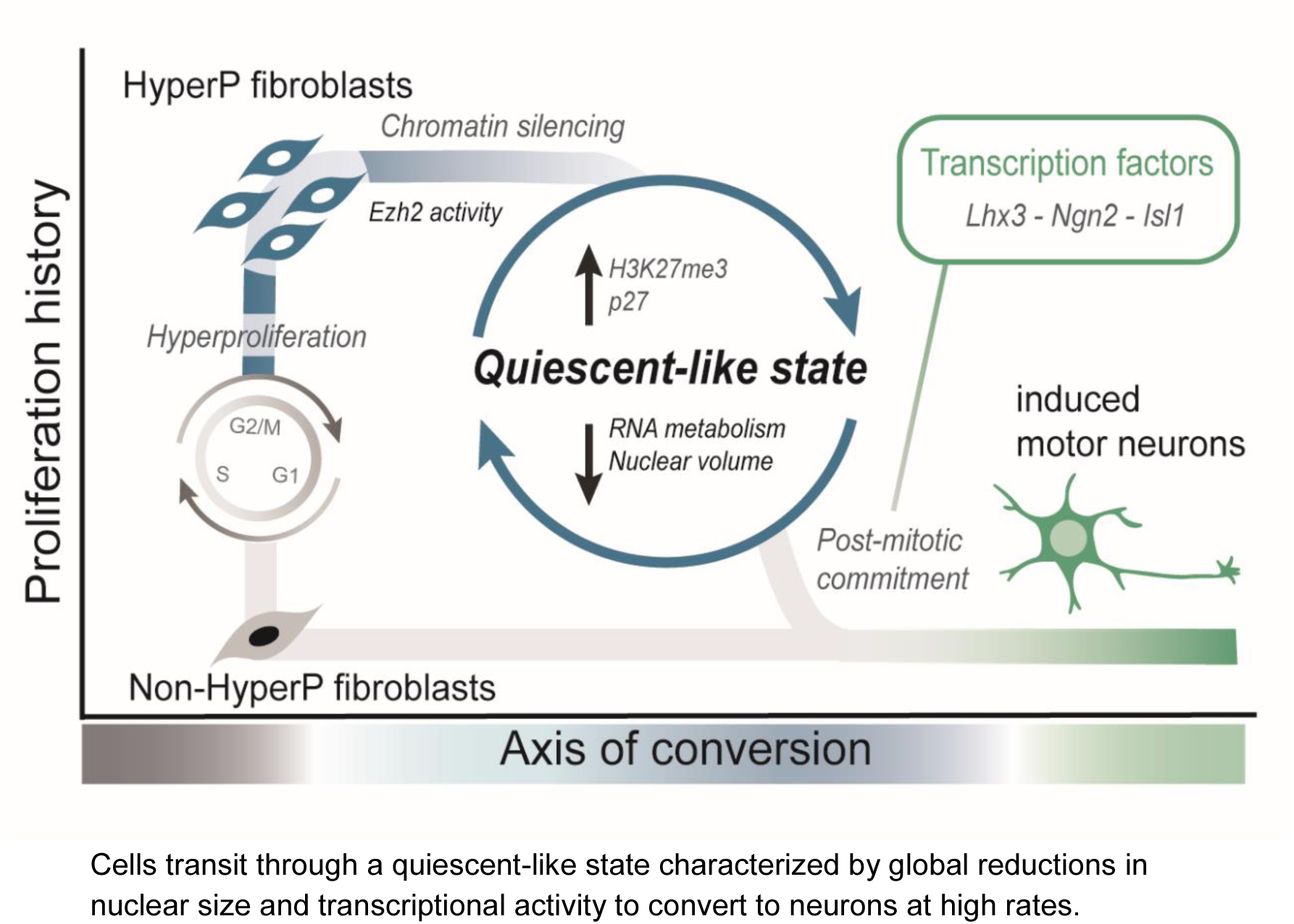

## Introduction

Transcription factors (TFs) provide essential cues for directing and redirecting cell fate in native and synthetic cell-fate transitions. However, the expression of lineage-specifying TFs alone infrequently drives somatic cells to adopt a new identity.^1–17^ In development, a synchronous set of events—including waves of transcription, cell division, and induction of TFs—guide progenitor cells through differentiation.^18–21^ The highly choreographed induction of genes suggests regulatory structures exist to reinforce the coordination of diverse processes that define the cell state.^22^ Loss of lineage-specific epigenetic and transcriptional programs can impede differentiation.^19^ Putatively, progenitors retain a state of plasticity through specific molecular mechanisms which may be common in other cell-fate transitions.^3,17,23–26^ Cells generate new fates by the activity of transcription factors. Yet how do some cells become receptive to transcription factors while others remain refractory? Processes that support cell-fate transitions are highly entangled, impeding simple inference of the features that confer plasticity.^3,23,27,28^ Potentially, comparison of sequences of events that co-occur in native and synthetic transitions may illuminate common motifs and mechanisms that confer receptivity.

Direct conversion of somatic cells to neurons offers a parallel system to examine neuronal fate commitment that is unconstrained by the coordinated processes of normal development. Thus, cell states that arise in both conversion and development may reveal common, possibly essential characteristics of this cell-fate transition. During conversion, cells do not necessarily transit through a known progenitor state^29,30^ and proliferation is not required for reprogramming.^29,31–33^ However, highly proliferative cells reprogram at higher rates to both iPSCs and post-mitotic neurons.^3,15–17^ Similarly, prior to differentiation, neural stem cells divide before neuronal fate commitment.^34^ In both cell fate transitions, proliferation may support commitment.

Potentially, proliferation drives cells to a state that is more receptive to the instructive cues from transcription factors and lineage-specific signaling.^28,35–39^ DNA replication limits active transcription and, at the same time, provides an opportunity for TFs to compete for access to the genome.^40^ In absence of competing factors, the symmetrical deposition of existing chromatin modifications allow propagation of chromatin states through recruitment of read-write enzymes.^40,41^ The division of chromatin modifications between daughter strands following replication may transiently dilute chromatin modifications and create a window of opportunity for TFs.^40–42^ TFs may therefore act in this window to compete with the propagation of histone marks over multiple cell cycles. Through passive loss of chromatin marks and by reducing the concentration of TFs^17^, proliferation may favor specific TFs that are retained on chromatin during mitosis.^43–46^ Thus, proliferation may support cell fate transitions by progressively priming the enhancer network and preparing for lineage-specific enhancer commissioning, while decommissioning enhancers of alternative fates.^46^

Direct conversion provides a well-defined model system to disentangle the intertwined role of proliferation in facilitating changes in cell fate from the cell fate itself.^15,17,29,47^ In particular, direct conversion of fibroblasts to motor neurons is well-defined with established murine transgenic reporters for staging the reprogramming process in living cells.^15,17,29,47^ Importantly, direct conversion to a post-mitotic identity provides several advantages over reprogramming to iPSCs. Conversion to a post-mitotic cell type, such as motor neurons, allows us to decouple transient changes in proliferation from proliferation associated with the new cell identity. Unlike transcription factors used to generate iPSCs, motor neuron transcription factors drive cells to exit the cell cycle.^48^ In other words, direct conversion to a post-mitotic cell type allows us to examine cells that are transiting *through* a proliferative state instead of transiting *to* a proliferative state. To improve our ability to track states that result in successful conversion, we have recently improved the robustness and efficiency of reprogramming to generate larger numbers of reprogrammed cells.^15,17^ Importantly, we developed live-cell labeling methods to isolate cells with different reprogramming potentials.^17^ This system allows us to finely dissect the molecular processes and cellular states that support conversion. With the direct conversion to neurons, we can identify transient states that would be nearly-inseparable during reprogramming to a highly proliferative cell type.

Here, using the model of direct conversion of mouse embryonic fibroblasts (MEFs) to induced motor neurons (iMNs), we identify a highly receptive cell state that emerges following hyperproliferation. Driven to this transient cell state by hyperproliferation, cells convert at high rates in response to the activity of neuronal transcription factors. By profiling patterns of chromatin accessibility and transcriptional activity in single cells, we find that cells transit to neurons via a quiescent-like state characterized by globally-elevated levels of H3K27me3, low transcriptional activity, reduced nuclear volume, and increased markers of quiescence. Proliferation induces global increases in H3K27me3, which supports cells’ transit through a state of altered RNA metabolism. Inhibition of Ezh2, the catalytic subunit of the Polycomb repressor complex 2 (PRC2) responsible for depositing a tri-methyl mark on H3K27, compromises the cells’ transit to this quiescent-like state, abolishing conversion to iMNs. Cells rapidly and simultaneously remodel their profile of biomolecules and epigenetic states, allowing transit through a quiescent-like state. Putatively, in this quiescent-like state, cells become receptive to neuronal transcription factors through the global weakening of interlocking forms of epigenetic and transcriptional feedback that reinforce fibroblast identity.

## Results

### Reduced accessibility across the axis of conversion defines reprogrammable cells

To induce direct conversion, we transduced transgenic primary mouse embryonic fibroblasts (MEFs) bearing the motor neuron transgenic reporter Hb9::GFP with retroviruses encoding motor neuron transcription factors Lhx3, Ngn2, and Isl1 (LNI) on a single transcript.^17^ The pioneer factor Ngn2 is essential for conversion to induced motor neurons (iMNs).^17^ Substitution of the pioneer factor Ngn2 with a DNA-binding domain^49^ mutant of Ngn2 abolishes the formation of iMNs (**Figure S1a-b**). For high efficiency conversion, we include a chemo-genetic cocktail consisting of two oncogenes, HRas^G12V^ and p53DD, encoded on a single transcript (RIDD) that are delivered virally, and the small-molecule TGF-β inhibitor RepSox (**Figure 1a)**.^17^

**Fig 1.**
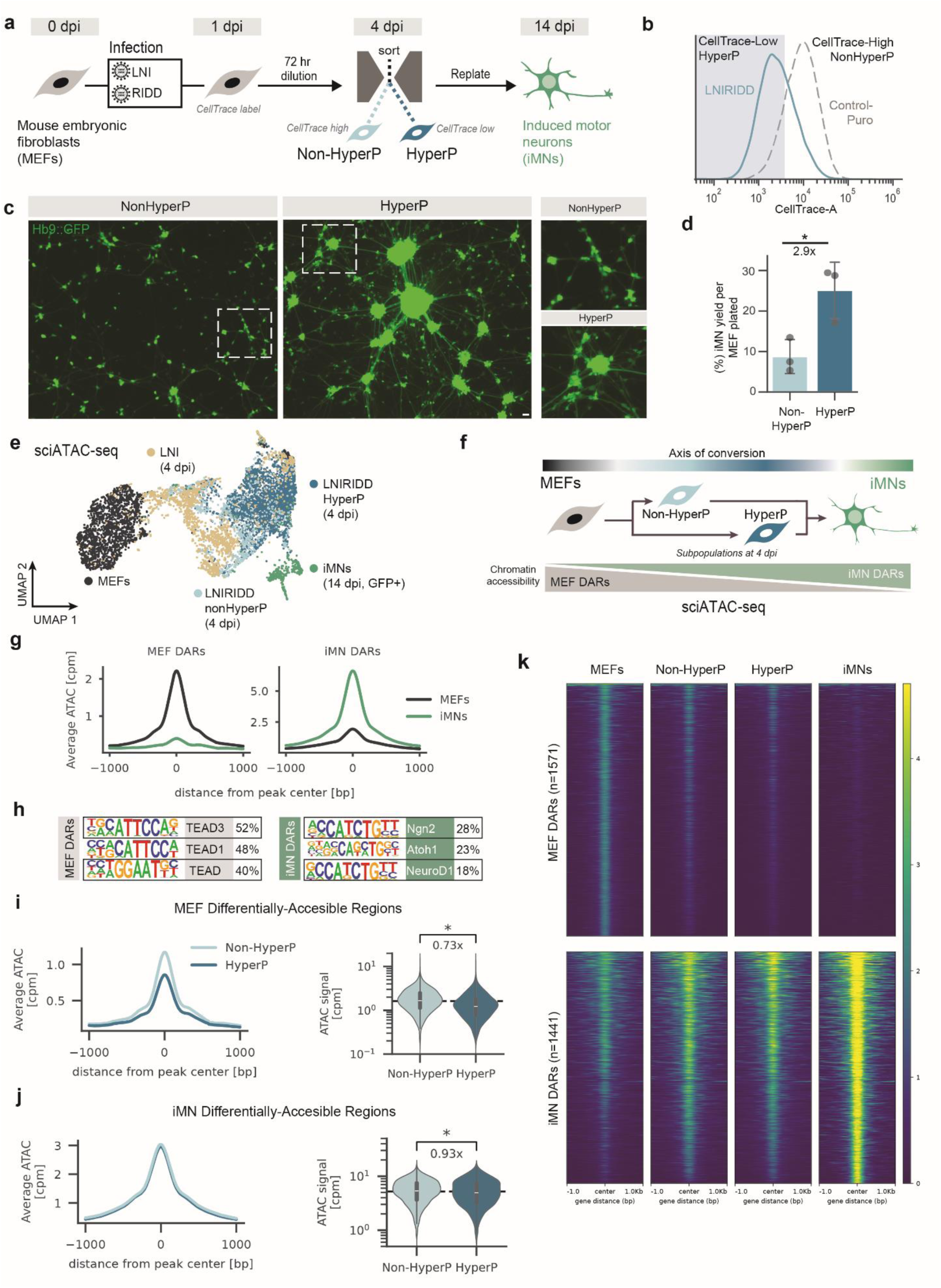
Reduced accessibility across the axis of conversion defines reprogrammable cells. a. Schematic for MEFs-to-iMN conversion via retroviral transduction of polycistronic cassette Lhx3, Ngn2, and Isl1 (LNI) supplemented with chemo-genetic cocktail HRas^G12V^-IRES-p53DD (RIDD) and RepSox. CellTrace dye dilution for 72 hours was used to identify subpopulations with history of hyperproliferation (HyperP) at 4 dpi. b. CellTrace distributions of Control-Puro cells and LNIRIDD-transduced cells. Hyperproliferative cells have CellTrace signal lower than the dimmest 20% of Control-Puro cells at 4 dpi (grey shaded region) c. Representative images of Hb9::GFP at 14 dpi for HyperP and Non-HyperP cells isolated using FACS and re-plated at 4 dpi, showing differences in morphology and conversion rate. d. iMN yield, the number of iMNs at 14 dpi divided by the number of cells re-plated at 4 dpi after FACS for HyperP and Non-HyperP cells quantified by flow cytometry. Bars represent the mean of three replicates. Dots represent the yield for an individual replicate. e. Uniform Manifold Approximation and Projection (UMAP) plot of single cell chromatin accessibility profiles. Each dot represents a single cell. f. During conversion, cells lose their fibroblast-associated chromatin profile by closing MEF-associated differentially-accessible regions (DARs) and adopt a motor neuron-associated chromatin profile by opening iMN-associated DARs. Cells can be staged along the “Axis of Conversion” by examining the relative closing of MEF DARs and opening of iMN DARs. g. Average read-depth normalized chromatin accessibility signal in 2000 base pair [bp] regions centered on the center of MEF (left) and iMN (right) DARs for MEFs and iMNs showing MEFs have greater accessibility in MEF DARs than iMNs and the opposite for iMN DARs. (cpm = counts per million NGS reads) h. Selected transcription factor motifs identified as enriched in MEF (left) and iMN (right) DARs and the fraction of peaks in the set of DARs that contain the motif shown. DARs shown are enriched motifs that are unique to either DAR set. i. Average read-depth normalized chromatin accessibility signal in MEF DARs for LNIRIDD HyperP and Non-HyperP cells (left). Violin plot showing distribution of chromatin accessibility signal in MEF DARs for the two populations. Data from high quality single cells were merged to create a bulk-accessibility profile. Statistical significance determined by one-tailed Kolmogorov–Smirnov test. Fold change indicated is fold change of the median values. j. Average read-depth normalized chromatin accessibility signal in iMN DARs for LNIRIDD HyperP and Non-HyperP cells (left). Violin plot showing distribution of chromatin accessibility signal in iMN DARs for the two populations. Data from high quality single cells were merged to create a bulk-accessibility profile. Statistical significance determined by one-tailed Kolmogorov–Smirnov test. Fold change indicated is fold change of the median values k. Heatmap showing the read depth normalized chromatin accessibility profiles for each individual DAR in the MEF DAR and iMN DAR sets for the conditions shown. Data from high quality single cells were merged to create a bulk-accessibility profile. Significance summary: p> 0.05 (ns); *p ≤ 0.05; ** p ≤ 0.01; ***p ≤0.001; Scale bars = 50µm. Fold changes are calculated based on Group means.

Cells with a history of hyperproliferation convert to neurons at higher rates.^17^ In direct conversion, neuronal transcription factors drive cells to a post-mitotic state.^15,17,47^ Thus, a history of hyperproliferation during direct conversion indicates that cells have rapidly divided before establishing a post-mitotic identity.^17^ To assess proliferation history, we label cells with a stable CellTrace dye at 1 dpi, allow the cells to divide and dilute the dye for 72 hours, and quantify CellTrace signal via flow cytometry at 4 dpi. We then use fluorescence-activated cell sorting (FACS) to isolate cells with a history of hyperproliferation (HyperP cells). HyperP cells have a lower CellTrace signal compared to cells that have not divided as frequently in this time window (Non-HyperP cells) (**Figure 1a**). We define HyperP cells as those cells with CellTrace signal dimmer than the dimmest 20 percent of MEFs infected only with a control retrovirus expressing the Puromycin-resistance gene (Control-Puro) (**Figure 1b)**. RIDD increases the percentage of HyperP cells in MEFs (**Figure S1c-d**). Motor neuron transcription factors alone do not induce proliferation (**Figure S1c-d**). As expected, conditions with a high fraction of HyperP cells show higher cell numbers at 4 dpi (**Figure S1e**). Activity of the motor neuron transcription factors is necessary for conversion. However, deletion of the DNA-binding domain of Ngn2 does not affect proliferation (**Figure S1f**).

At 4 dpi, we isolated the Non-HyperP and HyperP subpopulations of MEFs infected with LNI and RIDD viruses (LNIRIDD) by FACS and re-plated the subpopulations at equal seeding densities. Reprogramming outcomes for each subpopulation were tracked to 14 dpi (**Figure 1c)**. At 14 dpi, we quantified the number of Hb9::GFP positive cells via flow cytometry. To assess conversion, we calculated the iMN yield, defined as the number of Hb9::GFP positive cells divided by the number of cells re-plated at 4 dpi. HyperP cells convert at nearly three times the rate of Non-HyperP cells (**Figure 1d)**.^17^ HyperP cells have lower, not higher, levels of LNI transcription factors than Non-HyperP cells^17^. Thus, we could not attribute differences in conversion to increased levels of transcription factors. Rather, we hypothesize that proliferation induces a cell state that is more receptive to transcription factor-driven conversion.

Given the substantial differences in conversion potential, we considered that differences in chromatin state may render HyperP cells more receptive to transcription factors compared to non-HyperP cells. To examine chromatin state, we used single cell assay for transposase-accessible chromatin with combinatorial indexing and high-throughput sequencing (sciATAC-seq)^50^ to profile LNIRIDD HyperP and Non-HyperP cells and compare their accessibility profiles to the starting and final cell types, MEFs and iMNs, respectively (**Figure 1e**). In addition, we also profiled LNI-only cells.

To convert, cells open genomic regions that are specific to the motor neuron identity and close regions that are specific to fibroblasts, generating an axis of conversion. We defined the axis of conversion as regions that are differentially accessible between MEFs and iMNs (**Figure 1f**). We hypothesized that, given their higher rates of conversion, the HyperP subpopulation would adopt a chromatin state closer to iMNs and more distant from fibroblasts compared to non-HyperP cells. Alternatively, hyperproliferative cells may represent a state that is merely more receptive to transcription factors. If the features that support plasticity are orthogonal to the axis of conversion, cells with greater receptivity may not be distinguishable by differences in chromatin accessibility across this axis.

After filtering for high quality cells (**Figure S1e**), we identify differentially accessible regions (DARs) between MEFs and iMNs (**Figure 1g**). MEF and iMN DARs are enriched for transcription factor motifs associated with their respective cell types (**Figure 1h**), validating our approach for identifying representative regions across the axis of conversion. We find the differences in proliferation history are reflected in chromatin accessibility (**Figure 1i-k, Figure S1f-h**). Interestingly, HyperP cells do not have greater accessibility at iMN DARs than Non-HyperP cells. Instead, HyperP cells have decreased ATAC signal at MEF DARs compared to Non-HyperP cells. Thus, conversion potential more strongly correlates with closing of fibroblast-associated chromatin rather than opening regions associated with the motor neuron identity.

### Receptive cells display reduced nuclear volumes, increased silent euchromatin

Changes in accessibility and chromatin structure may reflect or be reflected by changes in nuclear compaction and morphology, including nuclear volume.^51–55^ To better define the connection between reprogramming efficiency, chromatin organization, and nuclear state, we explored whether transposition patterns correlate with broader nuclear characteristics. As nuclear condensation has been linked to hyperproliferation^56^, we hypothesized that highly proliferative cells would show increased chromatin compaction and reduced nuclear volume.

To observe nuclear morphology, we examined cells reprogramming with the transcription factors in low and high efficiency conditions. In the presence of RIDD, most cells are hyperproliferative (HyperP; 75%) whereas in the absence of RIDD, the majority of cells with LNI are non-hyperproliferative (Non-HyperP; 83%) (**Figure 2a-b**). Measuring *in situ* we can avoid the confounding influence of dissociation on our ability to observe nuclear features such as area and volume. We used confocal microscopy to capture z-stacks of nuclei at 4 dpi. By reconstructing these projections, we calculated the nuclear volume and obtained across-sectional area of the nucleus. Cells in the LNIRIDD condition displayed significantly smaller, more compact nuclei compared to cells in the LNI condition (**Figure 2c-e**). In highly proliferative conditions, cells showed nearly a two-fold reduction in nuclear volume and cross-sectional area (**Figure 2d-e**).

**Fig 2.**
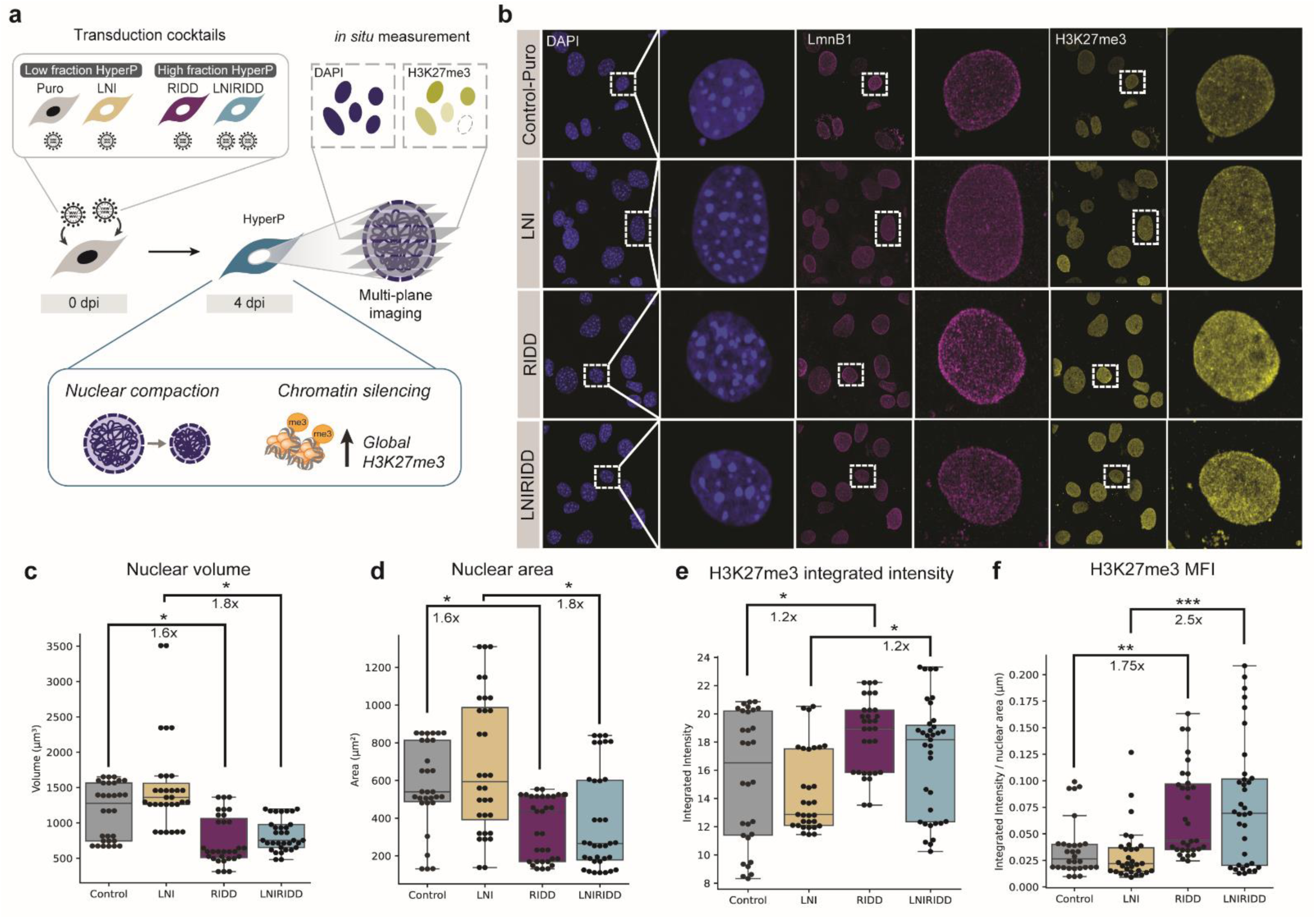
Receptive cells display reduced nuclear volumes, increases in silent euchromatin. a. Transduction cocktails generate different fractions of hyperproliferative cells at 4 dpi. Nuclear volume and H3K27me3 can be measured *in situ* using multi-plane confocal microscopy for these conditions and will broadly reflect differences in proliferative subpopulations. b. Representative images of cells fixed at 4 dpi and labeled with DAPI and stained for LmnB1 to delineate the nuclear lamina and H3K27me3. c. Nuclear volume of single cells determined by integration of z-stack areas using LmnB1 to define the end of the nucleus (n=23). d. Nuclear area of single cells determined as the largest area of from the stack of z-slices images. e. Integrated fluorescence intensity of H3K27me3 immunofluorescent staining for single cells. f. Mean fluorescence intensity of H3K27me3 immunofluorescent staining for single cells determined by dividing the integrated intensity by the nuclear area. Significance summary: p> 0.05 (ns); * p ≤ 0.05; ** p ≤ 0.01; ***p ≤0.001; Scale bars = 10µm. Fold changes are calculated based on Group means.

Changes in compaction and chromatin structure can affect accessibility of chromatin-binding proteins to their targets which may reflect specific histone modifications.^51^ We wondered if the substantial reduction in nuclear volume coincides with a global transition to a silent chromatin state marked H3K27me3. Staining for H3K27me3 reveals that LNIRIDD cells display elevated levels of total H3K27me3 than LNI-only cells (**Figure 2c, f-g, Figure S2a-b**). We do not observe an increase in H3K9me3 (**Figure S2a, c**). As nuclear area decreases, the mean intensity of H3K27me3 per nuclear area further increases (**Figure 2e, g**). In the presence of RIDD, cells show an increase in the levels of H3K27me3 per nuclear area (**Figure 2g**), suggesting a correlation between compaction and H3K27me3 in highly proliferative cells.

### Hyperproliferative cells broadly enrich H3K27me3 across the genome

Potentially, H3K27me3 may preferentially enrich in fibroblast-associated genomic regions, accelerating reprogramming by suppressing donor cell identity. Alternatively, H3K27me3 may globally increase across the genome, regulating both fibroblast and iMN gene regulatory networks. To examine these competing models, we used Cleavage Under Targets & Tagmentation (CUT&Tag)^57^ with spike-in normalization to identify regions marked by H3K27me3 in LNIRIDD HyperP and Non-HyperP cells as well as MEFs for reference (**Figure 3a**). Additionally, we profiled H3K4me3 to examine the distinct patterns of regulation across the genome.

**Figure 3.**
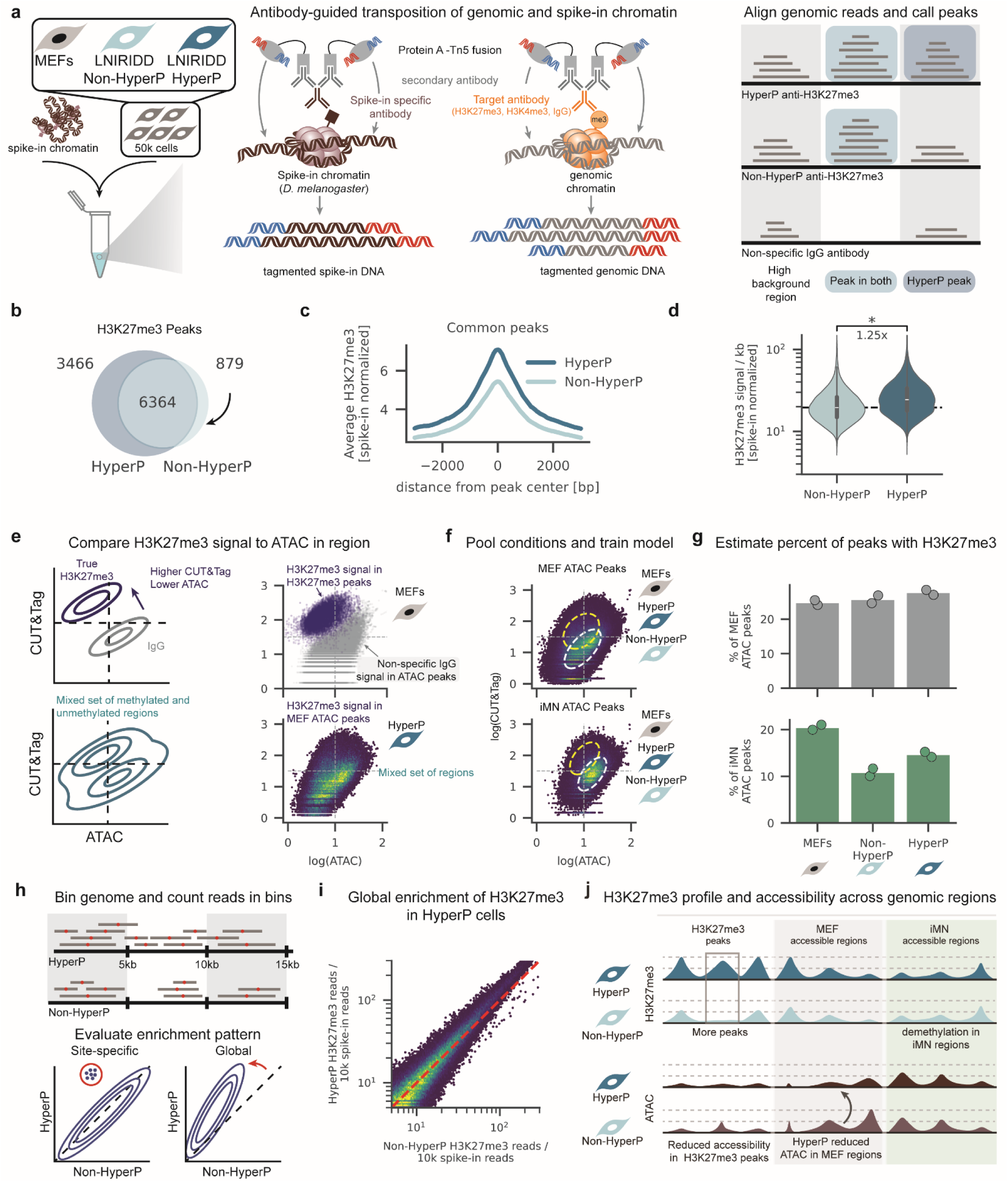
Hyperproliferative cells broadly enrich H3K27me3 across the genome. a. Schematic of Spike-in normalized CUT&Tag. A fixed amount of *D.melanogaster* chromatin with the H2Az histone are added to 50 thousand reprogramming cells FACS separated by proliferation history. After cell permeabilization, a spike-in antibody (anti-H2Az) and an antibody towards the protein of interest (anti-H3K27me3) are added, followed by a secondary antibody, and finally a Protein A-Tn5 transposase fusion protein that cleaves and tags (tagments) chromatin surrounding the protein of interest. These fragments are sequenced using next generation sequencing and aligned to the host genome. Genomic regions with a high number of reads specific to the antibody of interest are considered marked by the protein of interest (i.e. H3K27me3) and are called peaks. b. H3K27me3 peaks in master peak list by condition. For each condition, only peaks called in replicate datasets are retained. The replicated peak lists are then merged into a master peak list, where overlapping peaks are merged into a single contiguous region. c. For peaks called in both HyperP and Non-HyperP conditions, the average spike-in normalized coverage is plotted for regions spanning 3 kb in either direction from the peak center. d. H3K27me3 signal normalized by the number of spike-in reads and region length for peaks called in both HyperP and Non-HyperP conditions. Plot includes data from both biological replicates. Statistical significance determined by one-tailed Kolmogorov–Smirnov test. Fold change indicated is fold change of the median values. e. Expected correlation for a set of peaks that have H3K27me3 signal above background (top, left). “True” H3K27me3 should have high H3K27me3 signal and low ATAC signal while CUT&Tag signal from an experiment using a non-specific IgG antibody should correlate with accessibility. Scatter plot of H3K27me3 signal vs accessibility in H3K27me3 peaks (blue) in MEFs and IgG CUT&Tag signal vs accessibility in MEFs for regions defined as accessible in MEFs (MEF ATAC peaks) (top, right). When examining a pre-defined set of regions, the set may contain regions with only background CUT&Tag signal and regions with H3K27me3 signal above background (bottom, left). A mixed population is observed in the set of regions surrounding MEF ATAC peaks in HyperP cells (bottom, right) f. H3K27me3 CUT&Tag signal vs ATAC signal for ATAC peaks after pooling data from MEFs, LNIRIDD HyperP, and LNIRIDD Non-HyperP conditions used to train a Gaussian Mixture Model for the set of 10 kilobase regions centered on MEF (top) or iMN (bottom) ATAC peaks. The yellow ellipse corresponds to the Gaussian with higher H3K27me3 CUT&Tag signal than background. g. The fraction of each set of peaks estimated to be in the H3K27me3 high population (yellow ellipses in f) for MEF ATAC peaks (top) or iMN ATAC peaks (bottom). After training the Gaussian Mixture Model on the pooled data, the weight of the two Gaussians were re-optimized for each replicate for each condition to estimate the fraction of peaks within a single CUT&Tag dataset with high H3K27me3 signal. Bar represents the mean of the replicates, with individual replicates plotted as points h. Schematic of binning the genome into 5 kilobase (kb) regions and counting the number of reads in each bin (top) and the expected joint distribution for site-specific enrichment and global enrichment (bottom). Reads are assigned to a single bin by the location of the read midpoint (top, red dot). Global enrichment in one condition would manifest as a shift of the entire distribution above the x=y line (bottom, blue-dashed line). i. Joint distribution of H3K27me3 CUT&TAG signal in bins for hyperproliferative and non-hyperproliferative cells. Data for each condition is the mean, spike-in normalized read count of the two replicates. Red dashed line represents the x=y line. j. Global increases in H3K27me3 in HyperP cells manifests as an increase in the number of peaks a swell as the signal in the peaks over HyperP cells, but this global increase does not appear to be biased towards genomic regions associated with either the starting (MEF) or the final (iMN) cell state.

We first identified regions marked with H3K27me3 in hyperproliferative and non-hyperproliferative cells during conversion by calling peaks using MACS2^58^ and retaining peaks that are present in replicate datasets. H3K27me3 peaks called in our HyperP CUT&Tag dataset correlate with low accessibility in our HyperP sci-ATAC dataset, while H3K4me3 peaks correlate with high accessibility, validating our CUT&Tag assay and peak calling pipeline (**Figure S3a**). Annotating the locations of these peaks relative to annotated transcripts using HOMER^59^, we observe peaks called in hyperproliferative and non-hyperproliferative cells are distributed similarly across the genome (**Figure S3b**).

We combined the peak lists from hyperproliferative cells and non-hyperproliferative cells and merged overlapping peaks to create a master peak list with contiguous peaks. Of the genomic regions in this master peak list, a larger number are called peaks in hyperproliferative cells than in non-hyperproliferative cells (**Figure 3b**). We then quantified the CUT&Tag signal in these regions as the number of reads that overlap the region, normalized by the number of spike-in reads. Examining the common peaks between these two conditions, we find that the spike-in normalized H3K27me3 signal increases by ∼25% in hyperproliferative cells compared to non-hyperproliferative cells in these shared regions (**Figure 3c-d**). We also observed an increase in H3K4me3 signal in common peaks in HyperP cells (**Figure S3c**). As expected, H3K27me3 peaks are broader than H3K4 peaks (**Figure S3d**). H3K27me3 peak width distributions do not change significantly between hyperproliferative and non-hyperproliferative conditions (**Figure S3e**).

To erase donor identity and accelerate conversion, silencing may be focused at fibroblast regions. To examine this idea, we asked if H3K27me3 accumulated at or around regions defined as accessible in MEFs or iMNs (**Figure S4a**). Because of the low CUT&Tag signal at ATAC peaks (**Figure S4b**), we expect the data at ATAC peaks for each sample to contain subpopulations corresponding to background and true H3K27me3 signal (**Figure 3e**). The fraction of ATAC peaks falling in the “true” signal distribution corresponds to the fraction of ATAC peaks with enrichment of the H3K27me3 or H3K4me3 mark. To quantify the fraction of peaks with histone methylation above background, we fit a two-component, 2D Gaussian mixture model to the data. Specifically, we began by pooling data across all CUT&Tag samples for each histone mark and fitting a model for MEF ATAC peaks and iMN ATAC peaks separately (H3K27me3: **Figure 3f, Figure S4f**, H3K4me3: **Figure S4c-e, g**). By optimizing the weights of each Gaussian, the model provides an unbiased estimate of the fraction of peaks with histone mark enrichment over background within each sample. As estimated by our mixture model, the fraction of iMN ATAC peaks that have high H3K27me3 decreases between MEFs and both converting populations (**Figure 3g**). HyperP cells show a marginal increase in the fraction of MEF ATAC peaks with high H3K27me3. For both the iMN and MEF ATAC peak sets, the fraction of peaks with high H3K4me3 changed very little between populations (**Figure S4e**). Our data suggests that during conversion, iMN ATAC peaks lose H3K27me3, but proliferation does not specifically focus H3K27me3 increases within MEF-associated regions.

We wondered if proliferation drives a global increase in H3K27me3 levels. Expanding beyond predefined peaks, we sought to examine global changes in H3K27me3 levels. To do this, we binned the genome into 5 kb regions, assigned CUT&Tag reads to bins based on the midpoint of the read, normalized the binned counts using spike-in normalization, and compared the normalized signal between HyperP and Non-HyperP cells for signs of either site-specific or global enrichment (**Figure 3h**). When comparing signal across bins between replicate datasets, we observe strong correlation and bins are distributed around the x=y line (**Figure S4h-i**). When comparing signal between HyperP and Non-HyperP cells, we observe that the joint distribution shifts above the x=y line, indicating a global increase in H3K27me3 signal in HyperP cells (**Figure 3i**). Taken together, our data suggests that levels of H3K27me3 in hyperproliferative cells globally increase, gaining in intensity and number of peaks (**Figure 3j**). Changes in H3K27me3 and accessibility are not focused on specific regions corresponding to a specific cell identity. Rather, hyperproliferative cells show broad, genome-wide increases in H3K27me3 that coincides with reduced accessibility.

### Acute Ezh2 inhibition during induction abolishes iMN conversion

Given that high receptivity correlates with elevated levels of H3K27me3 at 4 dpi, we wondered if this mark serves a functional role in conversion. Alternatively, elevated levels of H3K27me3 may simply correlate with other processes that support receptivity. The polycomb repressive complex 2 (PRC2) deposits trimethyl marks on H3K27 via its catalytic subunit, Ezh2. In development, loss of polycomb can impede lineage priming and lead to stalled lineage commitment.^19^ Furthermore, loss of Ezh2 impedes reprogramming to iPSCs.^10^ Thus, we hypothesized that the ability to establish H3K27me3 may support receptivity by priming cells for transitions. To explore this hypothesis, we performed an acute inhibition of Ezh2 using a well-known small-molecule Ezh2 inhibitor, GSK126 (**Figure 4a**). Performing a dose titration, we identified a concentration for acute treatment that reduces H3K27me3 without inducing toxicity in MEFs (**Figure S5a-c**). Following acute treatment early in conversion (24 hrs, 1-2 dpi), we profiled reprogramming cells to observe changes in the fraction of hyperproliferative cells (**Figure 4b**), nuclear area, and levels of H3K27me3 at 4 dpi (**Figure 4c-e**). To examine the effect on reprogramming, we tracked conversion to iMNs at 14 dpi. At 4 dpi, GSK126 treatment slightly reduces the number of hyperproliferative cells (**Figure 4b**), suggesting a small effect on proliferation itself. As expected, inhibition of Ezh2 reduces levels of H3K27me3, generating a population with larger nuclei (**Figure 4d-f**).

**Fig 4.**
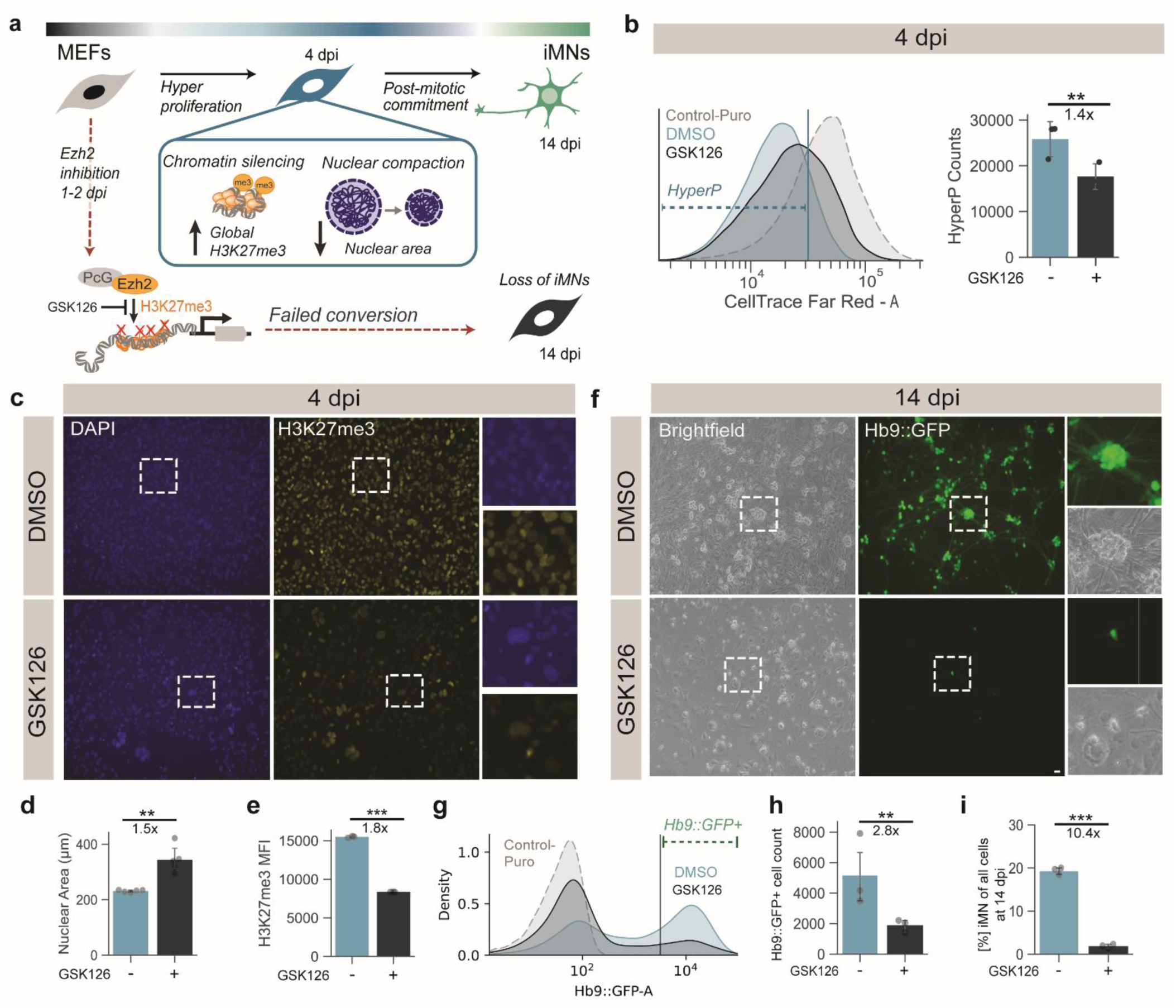
Acute Ezh2 inhibition during early induction reduces receptivity and abolishes iMN conversion. a. Conversion of MEFs to iMNs requires the activity of Ezh2. A transient inhibition of Ezh2 using GSK126 reduces levels of H3K27me3, increases nuclear area, and abolishes conversion. b. Proliferation of cells undergoing direct conversion with and without Ezh2 inhibition. Cells were treated with DMSO and 1 μM GSK126 for 24 hours from 1-2 dpi and quantified via flow cytometry at 4 dpi. (left) CellTrace distributions at 4dpi with DMSO or GSK126. (right) Number of cells at 4dpi classified as HyperP. c. Representative images of DMSO and GSK126-treated cells at 4 dpi stained with DAPI and immunostained for H3K27me3. d. Nuclear area at 4 dpi following DMSO or GSK126-inhibition from 1-2 dpi quantified via fluorescent microscopy. e. Mean fluorescence intensity of H3K27me3 immunofluorescence quantified via flow cytometry for LNIRIDD cells at 4dpi following DMSO or GSK126-inhibition from 1-2 dpi. Bar represent the mean of three replicates (dots). f. Representative images of conversion at 14 dpi. GSK126 inhibition reduces Hb9::GFP expression and adoption of neuronal morphology. g. Hb9::GFP fluorescence and iMN (Hb9::GFP+) gate for Puro-control (no fluorescence) and LNIRIDD with DMSO and GSK126 at 14dpi. gating strategy for quantification of Hb9::GFP-positive cells at 14 dpi. h. Quantification of total number of Hb9::GFP-positive cells at 14 dpi following DMSO or GSK126-inhibition from 1-2 dpi. i. Quantified purity of conversion. Purity is calculated by total number of Hb9::GFP-positive cells at 14 dpi over the total number of cells at 14 dpi. Significance summary: p> 0.05 (ns); *p ≤ 0.05; ** p ≤ 0.01; ***p ≤0.001; Scale bars = 10µm. Fold changes are calculated based on Group means.

Acute inhibition of Ezh2 abolishes conversion (**Figure 4f-i**). While a small number of Hb9::GFP-positive cells emerged, these cells displayed non-neuronal morphology, indicating incomplete conversion (**Figure 4f**). Additionally, inhibition of Ezh2 showed a selective vulnerability in reprogramming, reducing the total count of cells at 14 dpi (**Figure 4h, S5e**). Loss of H3K27me3 does not reduce cell counts in conditions with a low fraction of HyperP cells (**Figure S4d, e**), suggesting that H3K27me3 protects cells during periods of hyperproliferation. Together these data suggest that cells driven to proliferate enter a state that is selectively vulnerable to the loss of Ezh2 activity. Notably, cells that survive GSK126 treatment activate Hb9::GFP infrequently and do not establish proper neuronal morphology, indicating that the state of proliferation-driven receptivity relies on the cell’s ability to deposit H3K27me3 via Ezh2. Altogether we find that acute inhibition of Ezh2 prevents conversion and significantly reduces the fraction of hyperproliferative cells, leaving a population with larger nuclei and compromised receptivity to TF expression.

### H3K27me3 supports cells’ transit through a state of altered RNA metabolism

Given that transcription factor-mediated conversion relies on transcriptional activation, we wondered how global increases in marks of chromatin silencing might affect global transcription rates and RNA metabolism. We hypothesized that global increases of H3K27me3 would correspond to a global decrease in transcriptional activity (**Figure 5a**). Using 5-ethynyluridine (EU), we measured transcription rates by labelling nascent transcripts over one hour at 4 dpi.^15^ After fixing cells, transcripts were labeled by a “click” reaction to a fluorescent probe, which we visualized by microscopy (**Figure 5b**) and quantified via flow cytometry (**Figure 5c**). By co-staining for H3K27me3, we examined correlation between transcription rates and H3K27me3 signal with single-cell resolution. As we hypothesized, conditions with higher levels of H3K27me3 show lower rates of transcription, particularly the LNIRIDD condition (**Figure 5b-c, Fig S6a)**. Examining the joint distribution between EU and H3K27me3, we observe a decrease in EU signal and an increase in H3K27me3 levels (**Figure 5c**).

**Fig 5.**
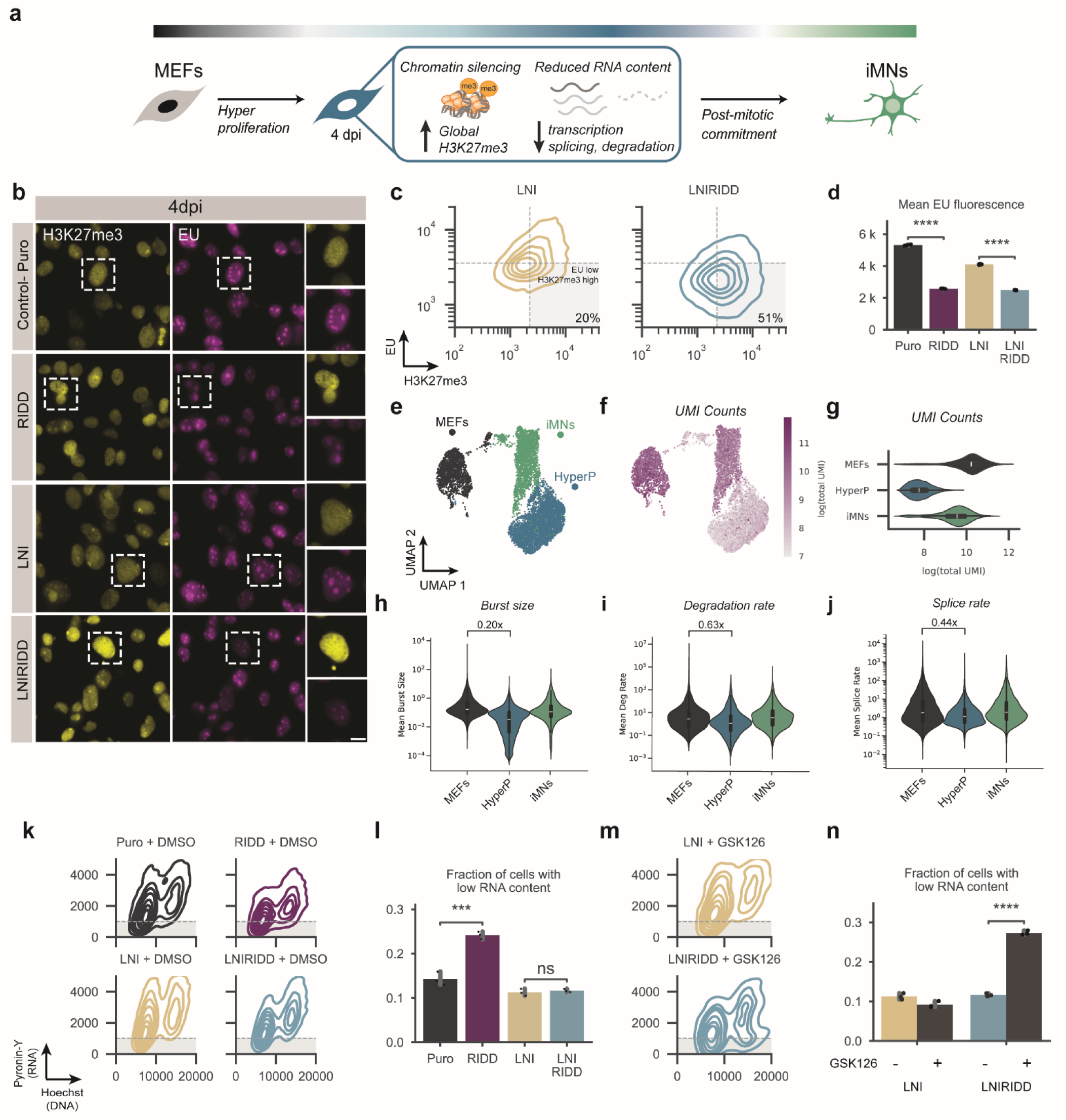
H3K27me3 supports cells’ transit through a state of altered RNA metabolism. a. During LNIRIDD-mediated reprogramming, hyperproliferative cells increase H3K27me3 and decrease RNA production. b. Representative fluorescent confocal microscopy images of 4 dpi cells stained for H3K27me3 and 5-ethynyluridine (EU), which was fed for 1 hour immediately preceding cell fixation and stained with azide-conjugated fluorophore. c. Kernel density estimate joint plot of cells stained for H3K27me3 and EU and quantified by flow cytometry for LNI and LNIRIDD at 4 dpi. d. Mean fluorescence intensity of EU signal measured by flow cytometry. Dots represent the mean fluorescence intensity of all cells from a single replicate, bars represent the mean of replicates (n=3). e. UMAP plot of scRNA-seq data for MEFs, 4 dpi LNIRIDD HyperP (HyperP), and iMNs from Wang et al. f. Log-transformed number of total transcripts (Unique Molecular Identifiers, UMIs) projected on to the UMAP. g. Distribution of log-transformed number of total transcripts per cell in the indicated condition. h. Distribution of burst sizes in the indicated condition as estimated by biVI. i. Distribution of RNA degradation rate in the indicated condition as estimated by biVI. j. Distribution of RNA splice rate in the indicated condition as estimated by biVI. k. Kernel density estimate joint plot of cells stained for Pyronin-Y and Hoechst to quantify DNA and RNA content at 4 dpi for the indicated condition with DMSO (vehicle control) treatment from 1-2 dpi. l. Fraction of cells with low RNA content at 4dpi, as defined by the grey shaded region in the joint plots in (k). m. Kernel density estimate joint plot of cells stained for Pyronin-Y and Hoechst to quantify DNA and RNA content at 4 dpi for the indicated condition with GSK126 treatment from 1-2 dpi. n. Fraction of cells with low RNA content at 4 dpi with and without GSK126 treatment. Statistical significance determined with student’s t-test. Significance summary: p> 0.05 (ns); *p ≤ 0.05; ** p ≤ 0.01; ***p ≤ 0.001; Scale bars = 5µm. Fold changes are calculated based on Group means.

We wondered if the reduced transcription rate might affect the number of coding transcripts. Lower transcription rates may translate to fewer poly-adenylated RNAs, which can be quantified in single cells using single-cell RNA-seq (scRNA-seq). Examining scRNA-seq data from Wang *et al.*^17^, we find that hyperproliferative converting cells show a substantial reduction in UMI counts per cell compared to both the starting MEF population and the final iMN population (**Figure 5e-g).**

To gain greater resolution on RNA metabolism, we turned to a recently developed computational tool, biVI, that can infer sizes of transcriptional bursts, rates of RNA degradation, and rates of RNA splicing.^60^ By integrating biophysical modeling, biVI uses transcript counts to infer rates of RNA metabolism from the distributions of nascent and mature RNA transcripts. Using biVI, we find that compared to MEFs and iMNs, hyperproliferative cells are estimated to have reduced burst sizes, rates of degradation, and rates of splicing, suggesting a global reduction in RNA metabolism (**Figure 5g-i).** The reduction in RNA burst sizes in HyperP cells concurs with the reduced transcription rates observed in EU-click assays (**Figure 5c**).

Cells with lower transcription rates may contain fewer total RNAs.^61^ To examine the total RNA content of the cells, we stained with Pyronin-Y following Hoechst labeling.^62^ Fluorescence signal increases from Pyronin-Y for increasing RNA content in single cells. We measured RNA and DNA content at 4 dpi using flow cytometry (**Figure 5k**). Overall distributions of cells across DNA and RNA content are mostly similar at 4 dpi across conditions (**Figure S6b**).

Given the changes in RNA metabolism, we wondered how inhibition of Ezh2 might affect RNA content. If Ezh2 supports transient silencing of chromatin, inhibition might generate a population of cells with higher transcription rates and more RNA content. Alternatively, if H3K27me3 supports the cells’ transit through a state of low RNA content, then cells may stall in this state, preventing the establishment of neuronal identity. At 4 dpi, cells treated with GSK126 show an altered distribution of RNA content (**Figure 5m, Figure S6b**). Inhibition of Ezh2 increases the fraction of cells with low RNA content, suggesting that H3K27me3 is not required to obtain a state of low RNA content (**Figure 5n**). Ezh2 inhibition and therefore loss of H3K27me3 generates a cell population with high DNA content but very low RNA levels, suggesting a “stalled” state. We had previously identified a sublethal concentration for GSK126 in MEFs (**Figure S5a-c**); however, in reprogramming, we notice that addition of RIDD with or without TFs generates a slight selective vulnerability to GSK126 treatment (**Figure S5d-e**). At 14 dpi, the surviving population of GSK126-treated cells is enriched for low RNA content (**Figure S6c**). Together, our data suggest that proliferation drives cells to a highly receptive state characterized by reduced RNA metabolism. In this receptive state, elevated levels of H3K27me3 protect cell viability and support cells’ transition through a state of low RNA content.

### Inhibition of Ezh2 reduces p27, blocks cells transit through a quiescent-like state

Adult neural stem cells emerge from quiescence to proliferate and differentiate into post-mitotic neurons.^34,63^ Neural stem cells are marked by expression of specific transcription factors that guard stem cell fate and poise cells for differentiation. Previous work suggests that fibroblasts do not convert to neurons through a definitive stem cell-like state marked by specific transcription factors.^29^ However, a quiescent-like state may provide a pivotal transient state for converting cells as they prepare to commit to a post-mitotic fate. Quiescent cells are marked by altered RNA metabolism, low transcriptional activity, reduced cell size, and elevated H3K27me3 and expression of p27Kip1 (p27).^64,65^ Potentially, cells may convert at high rates by transitioning through a quiescent-like state. PRC2-mediated increases in H3K27me3 provide a central mechanism for coordinating the entry and exit into quiescence.^66^ Thus, we hypothesized that loss of H3K27me3 may abolish conversion by preventing either entry into or exit from a quiescent-like state (**Figure 6a**).

**Fig 6.**
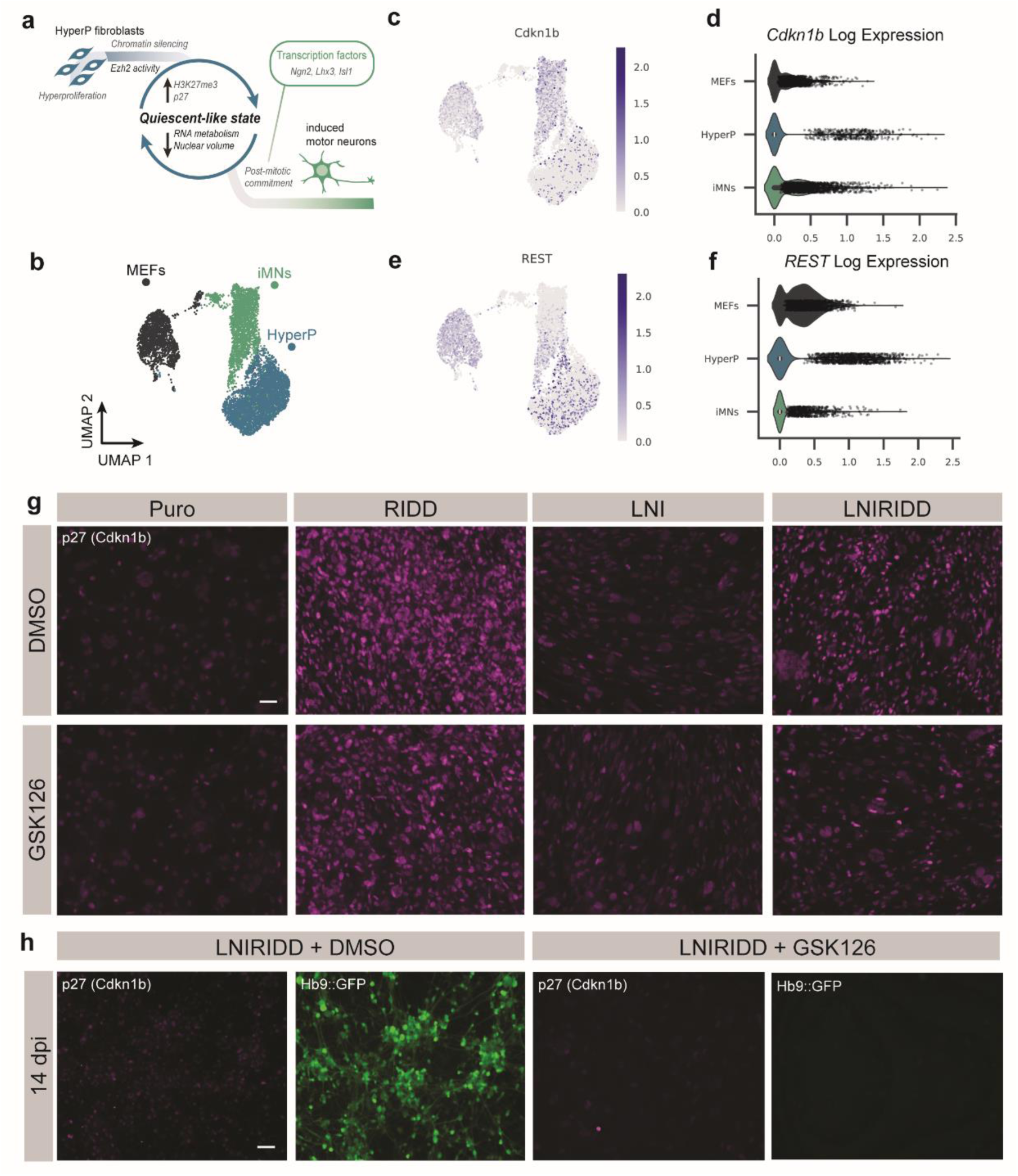
Inhibition of Ezh2 reduces p27, blocks cells transit through a quiescent-like state. a. Hyperproliferative cells attain a quiescent-like state marked by increased H3K27me3, decreased RNA metabolism, decreased nuclear volume, and increase expression of p27. Motor neuron transcription factors act on this state to drive cells into the post-mitotic iMN cell fate. b. UMAP plot of scRNA-seq data for MEFs, 4 dpi LNIRIDD HyperP (HyperP), and iMNs from Wang et al. c-d. CDKN1b (p27^Kip1^) is expressed in iMNs and a subset of HyperP cells. (c) log (Cdk1b) expression projected onto UMAP of scRNAseq data. (d) Distribution of log (Cdk1b) expression in each population of cells. Each dot represents expression of *Cdkn1b* from a single cell, and only cells with greater than 0 expression are plotted. e-f. REST (NSRF) is expressed in MEFs and a subset of HyperP cells. (e) log (REST) expression projected onto UMAP of scRNA-seq data. (f) Distribution of log (REST) expression in each population of cells Log expression of REST (NSRF) across reprogramming trajectory. Each dot represents expression of Cdkn1b from a single cell, and only cells with greater than 0 expression are plotted. g. Representative images of p27 immunofluorescent staining at 4 dpi in the indicated conditions. Scale bar = 50 µm. h. Representative images of p27 immunofluorescent staining and Hb9::GFP expression for LNIRIDD cells at 14 dpi without (left) and with (right) acute GSK126 treatment from 1-2 dpi. Scale bar = 50 µm.

We wondered if we could observe a transcriptional signature of a quiescent-like state. Examining scRNA-seq profiles, we observed cells with elevated expressions of two markers of quiescence, Cdkn1b (p27) and REST (**Figure 6b-e**). p27 marks quiescent cells and is implicated in both the entry and exit into the quiescent state.^67^ REST (also called the neural restrictive element-1 silencing transcription factor (NRSF)) restricts commitment to the neuronal fate in non-neuronal cells by blocking the binding of neuronal transcription factors to restrictive element-1 (RE-1) elements.^63^ REST restrains neurogenesis to maintain a population of quiescent stem cells.^63^ Both markers of quiescence were increased in a fraction of the hyperproliferative cells. We did not observe increases in progenitor markers (**Fig S6d**), suggesting they may not be expressed or are expressed below the limit of detection in scRNA-seq.

We were specifically intrigued by the high expression of Cdkn1b observed in a fraction of HyperP cells (**Figure 6c, d**).^64^ p27 supports differentiation from progenitors within the neuronal lineage and maintains postmitotic identity.^68^ High Cdkn1b expression is maintained in iMNs at 14 dpi, unlike other cell G1 and G0 markers including Cdk6, Rb and Dek (**Figure 6d; Fig S7a**). p27 is regulated post-translationally, affecting its stability and activity. Across most cells in the LNIRIDD condition, we observe high levels of p27 (**Figure 6g**).

Considering that p27 may mark or guide cells to neuronal fate, we wondered if inhibition of Ezh2 might impede transit of cells to or from a p27-positive state. Alternatively, Ezh2 inhibition may prevent conversion independent of p27. Staining for p27, we find that inhibition of Ezh2 and loss of H3K27me3 reduces the fraction of cells with high expression of p27 at 4 dpi (**Figure 6h, Figure S7b-c**). Loss of p27 coincides with absence of Hb9::GFP-positive cells at 4 dpi (**Figure 6h**). Moreover, as expected from the analysis of scRNA-seq data, iMNs at 14 dpi retained expression of p27 (**Figure 6d**).

## Discussion

Here, in examining fibroblast to motor neuron conversion, we have defined the features of cells that are highly receptive to cell fate programming via transcription factors. Hyperproliferation and increased conversion potential are associated with closing of donor cell-associated chromatin rather than opening final cell state-associated chromatin. We show that hyperproliferation drives cells to a state characterized by global reductions in nuclear size and reduced transcriptional activity. Supported by increased levels of H3K27me3, cells gain access to a quiescent-like state defined by reduced RNA metabolism and elevated expression of p27. Cells enter this quiescent-like state even in the absence of motor neuron transcription factors, suggesting these TFs are dispensable for entry into this state. Overexpression of the oncogenes with inhibition of TGF-β via RepSox drives hyperproliferation, routing cells to this quiescent-like state, dependent on the activity of Ezh2. Inclusion of the neuronal transcription factors allows cells to activate neuronal markers and transit to a post-mitotic state. Thus, we find that rapid proliferation drives cells to a quiescent-like state in which they are receptive to the activity of transcription factors which drive neuronal fate commitment.

While experimental techniques to examine cell fate transitions often highlight subsets of specific genes and loci, we find that global changes in cellular processes differentiate cells with different reprogramming potentials. Moreover, these differences not only mark these populations but appear essential in conferring receptivity to transcription factor-mediated conversion. Global synthesis rates normally scale with cell size.^69–73^ While we observe scaling, we also observe that rates vary dynamically and defy scaling during transitions. Could such global changes be relevant and observable in other cell fate transitions such as differentiation and malignant transformation?^61,64,74–82^ We hypothesize that these global reductions in RNA metabolism may generally facilitate cell fate transitions. However, due to their reduced RNA metabolism and RNA content, observing these subpopulations by transcriptional profiling requires particular consideration.^61,78,83,84^ Bulk transcriptional profiles primarily report on the state of cells with high RNA metabolism, obscuring hyperproliferative and quiescent-like cells within a mixed population. As hyperproliferative cells comprise a small fraction of cells in the absence of oncogenes, signatures of these cells would be difficult to capture by transcriptional profiling as they would contribute a small number of RNAs to the overall profile. While single cell transcriptional profiling could identify these subpopulations, analytical processes often exclude cells with low UMI counts, and normalization techniques may further bias identification and interpretation of expression in these cells.^77^ Our results suggest greater attention should be paid to measuring and controlling for global changes in transcription, proliferation, and other central metabolic processes.^77,85^

Cells retain memory through diverse biomolecules such as RNA, proteins and post-translational modifications, including those propagated in chromatin. Rapid proliferation accelerates the dilution of stable biomolecules, biasing cell fate.^3,15,86,87^ While many others have found that proliferation potentiates cell-fate transitions^1,3,7,9,16,88^, our system allows us to decouple proliferation from the activity of the fate-specifying transcription factors, as neuronal transcription factors do not promote proliferation.^11,15,17,30,48^ Decoupling the activity of transcription factors and proliferation allows us to ask how these two processes independently work to route cells into receptive states that support neuronal fate commitment. Questions remain on how proliferation and other features of cell cycle may erase inherited memories of cell identity.^3,9^ For instance, does the rate of division set the probability of conversion, or does each round of division bring the cell sequentially closer to reprogramming? The resolution of our proliferation assay and the heterogeneous nature of reprogramming primary cells poses a limitation to address these questions.

Prolonged perturbations in RNA homeostasis can drive cell fate changes.^82^ High rates of proliferation coincide with reduced nuclear volume and RNA metabolism. Reductions in the rate of transcription may accelerate the loss of transcriptional memory through two distinct mechanisms. As the transcription rate decreases, cells reduce the production of transcripts from the original gene regulatory network. Thus, lower rates of transcription lead to a smaller reservoir of transcripts that can reinforce the original cell fate. Models of mutually-exclusive two-state systems such as toggle switches offer analogous systems to understand cell fate switching.^89–91^ In a toggle switch, the transcription rate sets the size of the reservoir and thus the stability of the states.^92^ Depletion of the reservoir should destabilize cell states and make cells more receptive to cues that induce transitions to new states. Additionally, as RNAs can recruit transcription factors and chromatin regulators to sites of nascent transcription^93–96^, a reduced rate of transcription may compromise transcriptional memory mediated by transcription, supporting loss of donor identity.

While reduced RNA metabolism may hasten the loss of cell identity, reduced nuclear volume could accelerate the search by transcription factors to establish new sites of transcription.^97–99,99,100^ If transcription factor mobility is independent of nuclear volume, the search of the smaller volume require less time.^101^ Potentially, in a smaller nucleus, neuronal transcription factors can more readily find target sites to induce neuronal fate commitment. Globally-reduced transcriptional activity may direct common transcriptional resources to a smaller number of sites in the genome, weakening gene regulatory networks that reinforce the current cell state.^102,103^ Understanding how transcription factor mobility and engagement with the genome changes across cell states may refine our understanding of how transcription factor activity and nuclear volume change and adapt to support or impede cell fate transitions.^104–106^

While converting cells do not appear to transit through a progenitor state—as defined by the expression of progenitor transcription factors, the properties of our quiescent-like cells bear a striking resemblance to quiescent neural stem cells.^63,64^ Like quiescent stem cells, these cells bear high levels of H3K27me3 which may be essential to support cells’ transition through a state of altered RNA metabolism. Alternatively, high levels of H3K27me3 may canalize cell fate and prime cells for subsequent differentiation while preventing the precocious activity of neuronal transcription factors.^19,107,108^ However, what processes balance maintenance of the quiescent-like state and trigger differentiation? Early in conversion, cells rapidly proliferate in the presence of neuronal transcription factors, suggesting mitotic signals can transiently overcome the activity of neuronal transcription factors. However, successful conversion to motor neurons requires the activity of Ngn2, as deletion of the Ngn2 DNA binding domain abolishes reprogramming. While Ngn2 activity is necessary to drive cells from the quiescent-like state into neuronal fate commitment, hyperproliferative cells do not convert deterministically. Our previous work suggests that cells that establish high rates of transcription following rapid division convert at near-deterministic rates.^15^ Re-establishing higher rates of transcription may depend on a range of factors including the activity of the pioneer factor Ngn2. Potentially, H3K27me3 may prime gene regulatory networks for activation by a pioneer factor such as Ngn2. The activity of Ngn2 in the quiescent-like state will depend on other transcription factors, signals, and transcriptional machinery rather than solely on the levels of Ngn2.^17^ Future work exploring signaling cues in the quiescent-like state may reveal signals or combinations of signals that are necessary and sufficient to drive cells to neuronal fate commitment.

Overall, our work builds a model of dynamic regulation across scales and processes. Proliferation drives cells to a quiescent-like state characterized by global changes in nuclear features, chromatin state, and RNA metabolism. We hypothesize that global reduction of transcription serves to weaken two interlocking forms of feedback that reinforce the donor identity— the production of transcripts and transcriptionally-dependent chromatin states. The global changes in the quiescent-like state may prime the nucleus for the activity of transcription factors which drive neuronal fate commitment. While we’ve identified mechanisms that allow cells to convert to neurons, the quiescent-like state may be broadly receptive to the activity of diverse transcription factors and signaling cues which may direct cell fate in development, tissue homeostasis, and transformation.

## AUTHOR CONTRIBUTIONS

A.M.B. and J.T. performed reprogramming experiments, FACS-sorting, and ATAC-library generation. J.T. performed H3K27me3 CUT&Tag. A.M.B. performed the CUT&Tag analysis. C.J. generated the sci-ATAC library. A.M.B. and C.J. completed the sci-ATAC-seq alignment and analysis. J.T. performed microscopy and image-analysis with assistance from KI Microscopy Core Facility. N.B.W. generated the scRNAseq data, performed E.U. Click staining and imaging, Pyronin-Y flow cytometry analysis and Ngn2-delta-bHLH experiments. C.O. assisted in scRNAseq reprocessing and analysis. A.M.B., J.T., and K.E.G. wrote the manuscript. K.E.G. and J.K.B revised the manuscript and supervised the project.

## ACKNOWLEDGEMENTS

Research reported in this manuscript was supported by the National Institute of General Medical Sciences of the National Institutes of Health under award number R35-GM143033, by the National Science Foundation under the NSF-CAREER under award number 2339986, and the with funding from Institute for Collaborative Biotechnologies. N.B.W. is supported by the National Science Foundation Graduate Research Fellowship Program under grant No. 1745302. A.M.B. is supported by the National Cancer Institute under award number F99CA284280. J.T. is supported by the Novo Nordisk Foundation Postdoc Fellowship for Research Abroad grant number NNF22OC0073452. We thank the Koch Institute’s RA. Swanson (1969) Biotechnology Center (National Cancer Institute Grant P30-CA14051) for technical support, specifically the Flow Cytometry Core, Microscopy and the BioMicro Sequencing Core Facilities.

Josh Brickman was funded by the Lundbeck Foundation (R370-2021-617, R198-2015-412, R400-2022-769, and R286-2018-1534), the Independent Research Fund Denmark (DFF-8020-00100B, DFF-0134-00022B, and DFF-2034-00025B), the Danish National Research Foundation (DNRF116), the Novo Nordisk Foundation (NNF210C0070898), and the European Union (ERC, SENCE, AdG 101097979). The Novo Nordisk Foundation Center for Stem Cell Medicine (reNEW) is supported by the Novo Nordisk Foundation grant number NNF21CC0073729 and previously NNF17CC0027852.

We thank Brittany Lende-Dorn, Sneha Kabaria, Kasey Love, and Deon Ploessl for feedback on the development of the manuscript.

## DATA AND MATERIALS AVAILABILITY

Raw FASTQs files and aligned BAM files are deposited in SRA (Temporary Accession # 14561323). Raw FASTQs files and aligned BAM files are deposited in SRA (Temporary Accession # SUB14561323). Raw data and computer codes used for data analysis are available from the corresponding author.

## MATERIALS AND METHODS

### Cell lines and culture

Plat-E and mouse embryonic fibroblasts were cultured in Dulbecco’s Modified Eagle Medium (DMEM) with 10% fetal bovine serum (FBS) at 37 °C, 5% CO2. Plat-E cells were also supplemented 10 μg/mL blastocidin and 1 μg/mL puromycin every 3 passages.

### Mouse embryonic fibroblast isolation

Mouse embryonic fibroblasts (MEFs) were isolated from embryos with Hb9::GFP background between E12.5 to E14.5). Hb9::GFP+ embryos were identified be green fluorescence of the spinal column under blue light. After removing the head and internal organs to remove competing cells, embryonic tissue was enzymatically dissociated using 0.25% Trypsin-EDTA and physically dissociated with razors for 5 minutes. Trypsin was quenched using DMEM+10%FBS and the dissociated tissue was collected, pelleted and resuspended in 0.25% Trypsin-EDTA and triturated. Again, the tissue was quenched and pelleted. The tissue was resuspended in DMEM + 10% FBS, filtered through a 40 μm cell strainer, and plated on to 10 cm cell culture dishes coated with 0.1% gelatin (1 embryo per dish). The isolated MEFs were cultured and expanded in Dulbecco’s Modified Eagle Medium (DMEM) supplemented with 10% fetal bovine serum (FBS) in 5% CO2. MEFs were passaged at least twice before reprogramming experiments. All MEFs were tested for mycoplasma after isolation and confirmed negative.

### Reprogramming to Induced Motor Neurons (iMN) via retroviral transduction

Retroviruses encoding motor neuron transcription factors (LNI), the high-efficiency cocktail (RIDD), or the puromycin-resistance gene (Puro), were produced using the Plat-E retroviral packaging cell line. Plat-E cells were seeded in 6-well plates to obtain ∼80% confluency the following day, 700-850k cells per well. The following day, cells were transfected using linear polyethylenimine with 1.8 μg per well of the 6-well plate using a 4:1 mass ratio of polyethylenimine:DNA. The following day, the media in each transfected well was removed and replaced with 1.25 mL of DMEM + 10% FBS + 25 mM HEPES buffer. MEFs were seeded into 96-well plates at a density of 10k cell per well of a 96-well. This cell density per culture area was maintained when scaling up reprogramming experiments from 96-well plates. The next day, viral supernatant was collected, filtered through a 0.45 μm filter, and 1.25 mL fresh DMEM + 10% FBS + 25 mM HEPE was added again to the Plat-E cells. Transduction cocktails were mixed to contain 11 μL of each virus and 5 μg/mL polybrene and then supplemented with DMEM + 10%FBS to 100 μL per 96-well to be transduced. The media was removed from the cells and the cells were incubated with the transduction cocktail overnight. Transduction was repeated for a second day. One day after the second viral transduction was designated 1 dpi at which time the transduction cocktail was removed and replaced with 100 μL per 96-well fresh media. At 3 dpi, media was switched to N3 media (DMEM/F-12 supplemented with N2, B27, and neurotrophic growth factors, BDNF, CNTF, GDNF, FGF all at 10 ng/mL). For conditions with RIDD, N3 media at 3 dpi was also supplemented with 7.5 μM RepSox. Cells were maintained in N3 media until the experimental endpoint. For cells quantified via flow cytometry at 4 dpi, cells were dissociated using 0.25% Trypsin-EDTA. For cells quantified via flow cytometry at 14 dpi, cells were dissociated using DNase/papain. iMNs were defined by gating the brightest Hb9::GFP fluorescent cells as we have reported previously.^15,17^

### CellTrace Proliferation Assay and FACS Sorting

For proliferation assays, transduced cells were labeled with CellTrace™ Far Red or CellTrace™ Violet Cell Proliferation Kit at 1 dpi. Cells were incubated in CT-FarRed or CT-Violet diluted in PBS according to manufacturer’s specifications and incubated with dye for 30 minutes at 37°C at 1 day post-infection (1dpi).

Cells were then washed with FBS-containing media and returned to culture for 72-hours. CellTrace intensity was quantified via flow cytometry at 4dpi. Hyperproliferative cells were isolated by fluorescence-activated cell sorting using a BD Aria or a Sony MA900 cell sorter. For reprogramming experiments, sorted cells were re-plated at a density of 10k cells per well of 96-well plate immediately after sorting and cultured in N3 media supplemented with penicillin-streptomycin.

### Immunocytochemistry

Cells were plated on a glass-bottom plate and cultured until their experimental endpoint. Cells were then fixed with 4% paraformaldehyde (PFA), permeabilized and blocked with a solution containing bovine serum albumin (BSA) and Triton X-100. Cells were washed three times, and incubated with primary antibody overnight at 4 °C. Cells were washed again before adding corresponding fluorophore-conjugated secondary antibodies. Nuclei were counterstained with DAPI before image acquisition using fluorescence (Keyence) or confocal (Leica Sp8) microscope.

### Confocal microscopy and 3D rendering of nucleus

Confocal images of DAPI-stained cells were imaged using Leica Sp8 confocal microscope with appropriate laser excitation and emission settings. Z-stacks ranging from 20-60 steps, with 0.30 Step size were obtained to capture end-to-end layers of multiple nuclei. A minimum of 3 random fields from 3 separate wells were obtained. 3D Nuclear reconstruction and 3D volume measurements were performed using Imaris image analysis software.

### CellProfiler Analysis of Staining intensity

Nuclear staining intensity for DAPI, H3K27me3, and H3K9me3 was analyzed using CellProfiler.^105^ Images were imported into the software for nuclear segmentation using the IdentifyPrimaryObjects module. Segmented nuclei were verified by eye, before staining intensity quantification using the MeasureObjectIntensity module. Thresholds for object identification and intensity measurement settings were kept consistent across all samples.

### EU-Click Assay

To assess global nascent transcription, cells were labeled with 1mM of 5-ethynyl uridine (EU) for 1 hour before fixation. For microscopy, cells were processed using the Click-iT™ RNA Alexa Fluor™ 594 Imaging Kit (ThermoFisher, #C10330). Briefly, cells were fixed with 3.7% PFA, permeabilized with 0.1% Triton X-100, and EU was conjugated with a Alexa Fluor™ 594 azide with a copper-catalyzed click reaction. Cells were then immunostained for H3K27me3 using an Alex Fluor 647™ conjugated antibody (Cell Signaling Technology #12158). For flow cytometry, cells were first dissociated using 0.25% Trypsin-EDTA. Cells were pelleted and resuspended in 3.7% PFA for fixation. Cells were then permeabilized with 0.5% Tween-20 and EU was conjugated with a sulfo-Cy3.5-azide (Lumiprobe #2330). Cells were then stained for H3K27me3 before quantification.

### GSK126 inhibition in reprogramming cells

A dose-response test for GSK126 (Ambeed) was established by treating MEFs with varying GSK concentrations and assessing cell viability by LIVE/DEAD NIR and H3K27me3 intensity. Reprogramming cells at 1dpi were treated with 1 μM GSK126 or DMSO for 24 hours. The effect of GSK126 inhibition on nuclear compaction, H3K27me3, and hyperproliferation (HyperP) was quantified at 4 dpi using immunocytochemistry and flow cytometry, respectively. Moreover, the effect of GSK126 inhibition on overall reprogramming yield of induced motor neurons (iMNs) was quantified at 14 dpi by flow cytometry.

### RNA profiling by Pyronin-Y

To simultaneously measure RNA and DNA content, cells were stained with Pyronin Y and Hoechst and quantified by flow cytometry, as in Eddoudi et al 2017.^106^ Briefly, cells were dissociated using Trypsin and resuspended in 1 mL of appropriate cell culture medium with 20 μM Hoechst 33342 (Invitrogen) and incubated for 45 minutes at 37 °C. After 45 minutes, Pyronin Y (aablocks) from a 10 mg/mL stock solution was added to the cells to a final concentration of 1 μg/mL and incubated for 15 minutes at 37 °C. After 15 minutes, cells were transferred to ice and analyzed by flow cytometry.

### CUT&Tag Library Generation

CUT&Tag libraries for H3K27me3 and H3K4me3 were generated MEFs and FACS-isolated LNIRIDD HyperP and Non-HyperP cells at 4dpi according to the procedure in Kaya-Okur et al. 2019^55^ using the CUT&Tag assay kit (Cell Signaling Technologies), with modifications to include spike-in chromatin. 50,000 cells were resuspended in 100 uL and bound to Concanavalin A beads, to which 10 ng of Spike-In Chromatin from Schneider’s *Drosophila* Line 2 (S2) (Active Motif) was added. Cells were incubated with an H3K4me3, H3K27me3, or IgG negative control antibody as well as a spike-in specific antibody (anti-H2Av) for 1 hour at room temperature (all rabbit IgG antibodies). Cells were washed and then incubated with an anti-rabbit secondary antibody for 30 minutes. Cells were washed and incubated with Protein A-Tn5 fusion protein loaded with adapters in a high salt buffer lacking MgCl_2_. After washing thoroughly using to remove unbound Tn5, cells were incubated in a Tagmentation buffer containing MgCl_2_ for 1 hour at 37 °C to initiate Tn5 transposition at antibody-bound genomic locations. Transposition was stopped by the addition of a Stop Buffer containing SDS and Proteinase K. Cell were incubated at 58 °C to release tagmented chromatin, which was then purified and libraries were prepared for sequencing. Sequencing was performed on a Singular G4 instrument with 2×150 bp paired end reads.

### CUT&Tag Data Processing and Analysis

Raw sequencing data obtained from bulk CUT&Tag experiments were processed using standard bioinformatics pipelines. Reads were trimmed to remove adapters using trim-galore. Trimmed reads were aligned to the mouse GRCm39 genome using Bowtie2 using options “-X 700--local--very-sensitive-I 10--no-mixed--no-unal--no-discordant” and to the *Drosophila* dm6 genome using options”-X 700--local--very-sensitive-I 10--no-mixed--no-unal--no-discordant--no-overlap--no-dovetail.” Mapped reads were filtered with samtools (quality threshold 2, removing unmapped reads). Filtered reads were converted to a bed file with the location of each fragment spanning the paired end reads using bedtools *bamtobed* function using the “-bedpe” option and selecting the 1^st^, 2^nd^, and 6^th^ column of this output file. This bed file was used as input to the MACS2 peak caller with options “-f BEDPE-B--broad--nolambda--keep-dup all-q 0.01.” broadPeak outputs from MACS2 for two replicate datasets were merged into a single peak list with overlapping peaks merged into a single region. This peak list was then filtered to retain only peaks that overlapped peaks in both replicate datasets. For examining CUT&Tag signal in CUT&TAG or ATAC peaks, reads per region were counted using bedtools *coverage* for each replicate CUT&TAG dataset and normalized by the number of spike-in reads (reads per 10 thousand spike-in reads). For examining per base coverage, CUT&TAG data were converted to BigWig coverage files using bedtools *genomecov* and the UCSC *bedGraphToBigWig* functions. The deeptools package was used for generating the average profile plots and heatmaps.

### Single cell RNA-seq (scRNA-seq) Reprocessing and Analysis

scRNAseq (10X) data from Wang et al^17^, was reprocessed to reflect MEF, HyperP and iMN conditions. Sequencing data were aligned to a custom reference based on GRCm39 with the GFP marker gene added using CellRanger to generate cells-by-gene count matrices. Scanpy (v 1.10.3) was used to perform basic quality control/preprocessing, including normalization, removal of cells with fewer than 1000 counts, removal of genes expressed in fewer than 3 cells, and identification of the 2000 most highly variable genes. Scanpy was then used to generate the UMAP and quantify expression of the specified genes.

For biVI analysis scRNAseq data were aligned to the GRCm39 mouse referencfe genome with the *kb-python* (v 0.28.2) software package.^107^ Count matrices generated with the *nac* workflow distinguished spliced and unspliced counts for further analysis. Scanpy was used to perform basic quality control/preprocessing, including normalization, removal of cells with fewer than 1000 counts, removal of cells whose counts were greater than 30 percent mitochondrial genes, and identification of the 2000 most highly variable genes. RNA metabolism metrics were estimated for each condition using the biVI software package^60^ ( with a “Bursty” model of transcription and dispersion factor of “gene-cell.”

### Ngn2 – delta bHLH reprogramming

The Ngn2^ΔbHLH^ mutant was generated by deleting the basic-helix-loop-helix (bHLH) domain from Ngn2. Wildtype and mutant Ngn2 were then cloned in frame with a 2A-mRuby2 so that mRuby2 fluorescence could be used as a proxy for Ngn2 expression. A separate polycistronic Isl1-2A-Lhx3 was also included to induce motor neuron reprogramming, as well as the single transcript RIDD (hRasG12V-IRES-p53DD). Cassettes were cloned using Golden Gate cloning via BsaI into a pENTR entry vector and Sanger sequenced confirmed. Then cassettes were transferred into the retroviral pMXs-gw destination vector using Gateway cloning by LR clonase and whole plasmid sequenced. Reprogramming was done as normal using Plat-E retroviral packaging to deliver: 1.) the wildtype (Ngn2-2A-mRuby2) or mutant (Ngn2^ΔbHLH^-2A-mRuby2) Ngn2 variants 2.) Isl1-2A-Lhx3, and 3.) RIDD. Flow cytometry was used to quantify proliferation at 4 dpi using CellTrace-Violet staining as described earlier, and to quantify reprogramming at 14 dpi.

### sci-ATAC-seq assay and analysis

A sci-ATAC-seq library was prepared following the protocol outlined in Del Priore et al 2021^108^. Briefly, unloaded Tn5 (2 mg/mL; Diagenode C01070010-20) was loaded with barcoded transposition primers and an initial transposition was carried out to validate the Tn5 concentration per cell required for transposition to yield fragment distributions with a mean fragment size around 800-1000bp. The validated concentration used was 1 uL of 40-fold diluted Tn5 per well (1000 cells).

Five cell populations were sorted: 1) MEFs, 2) LNI treated cell at 4 dpi, 3) LNI-RIDD treated cells at 4dpi, sorted as HyperP, 4) LNI-RIDD treated cells at 4dpi, sorted as “not HyperP” (e.g. above the HyperP gate), 5) Hb9:GFP+ iMNs at 14dpi. These cells were fixed with a final concentration of 0.1% formaldehyde at room temperature for five minutes. Sixteen thousand (16k) cells were distributed onto the Tn5-barcoded transposition plate, with 1k cells per well. After split-pooling, the cells were distributed across eight PCR plates. The library was quantified using an Agilent Bioanalyzer.

The resulting library was sequenced using Singular G4, using a 2×150 run with 36bp index reads. Reads were mapped to the GRCm39 genome build using Bowtie2. Single-cell and condition barcodes were recognized using custom Python scripts, allowing single-base-pair mismatches in the variable and constant barcode regions. Mapped reads were filtered with samtools (quality threshold 30, removing unmapped reads), and deduplicated with Picard. Reads were by +4 bp on the (+) strand and-5 bp on the negative stand to account for (-) the offset between Tn5 binding and adapter insertion^109^. Reads for each condition were then stored in a 5-column fragment file containing chromosome name, paired-end fragment start coordinate, end coordinate, cell barcode, and the number of times this read is duplicated (1 for all reads in our pipeline after deduplication).

Fragment files were imported into SnapATAC2^110^ and cells were filtered to retain only the subpopulation of high-quality cells with a high number of fragments per cell and a transcription start site enrichment score >3. The minimum number of fragments per cell was set on a condition-specific basis to ensure all high quality cells were retained. Fragments from high quality cells were then split into two fragments centered on the fragment ends (the transposition sites) and extended 100 bp in each direction. These transposition-centered fragments were used as inputs for the MACS2 peak caller run with options “--nomodel –nolambda--keep-dup all-f BEDPE--call-summits-q 0.01” to call peaks for each condition. Shifting reads to transposition sites before MACS2 is equivalent to using the MACS2 options “---shift-100--extsize 200.” The transposition-centered fragments from each condition were also evenly split into 2 “pseudoreplicate” files and peaks were called on each pseudoreplicate. Self-consistent peaks were selected using the IDR package,^111^ retaining peaks with an IDR value of <0.05. Peak summits called from each condition were extended 150 bp in each direction (roughly the size of a nucleosome, 301 bp total region). Peaks from each condition merged into a single list and then sorted in decreasing order of their significance scores. Keeping the most significant peak, overlapping peak windows that had lower significance scores were identified and then removed, leaving a list of the most significant (by q-value), non-overlapping peaks. Differentially accessible regions were then identified using SnapATAC2’s *diff_test* function using this peak set as input, and retaining regions with an adjusted p-value <0.01.

For examining ATAC signal in CUT&TAG or ATAC peaks, transposition-centered fragments per region were counted using bedtools *coverage* for each ATAC condition and normalized by the read depth (counts per million transposition-centered fragments). For examining per base coverage, transposition-centered fragments used to generate BigWig coverage files using bedtools *genomecov* and the UCSC *bedGraphToBigWig* functions. The deeptools package was used for generating the average profile plots and heatmaps.

