## Supplementary Figures for "Cells transit through a quiescent-like state to convert to neurons at high rates"

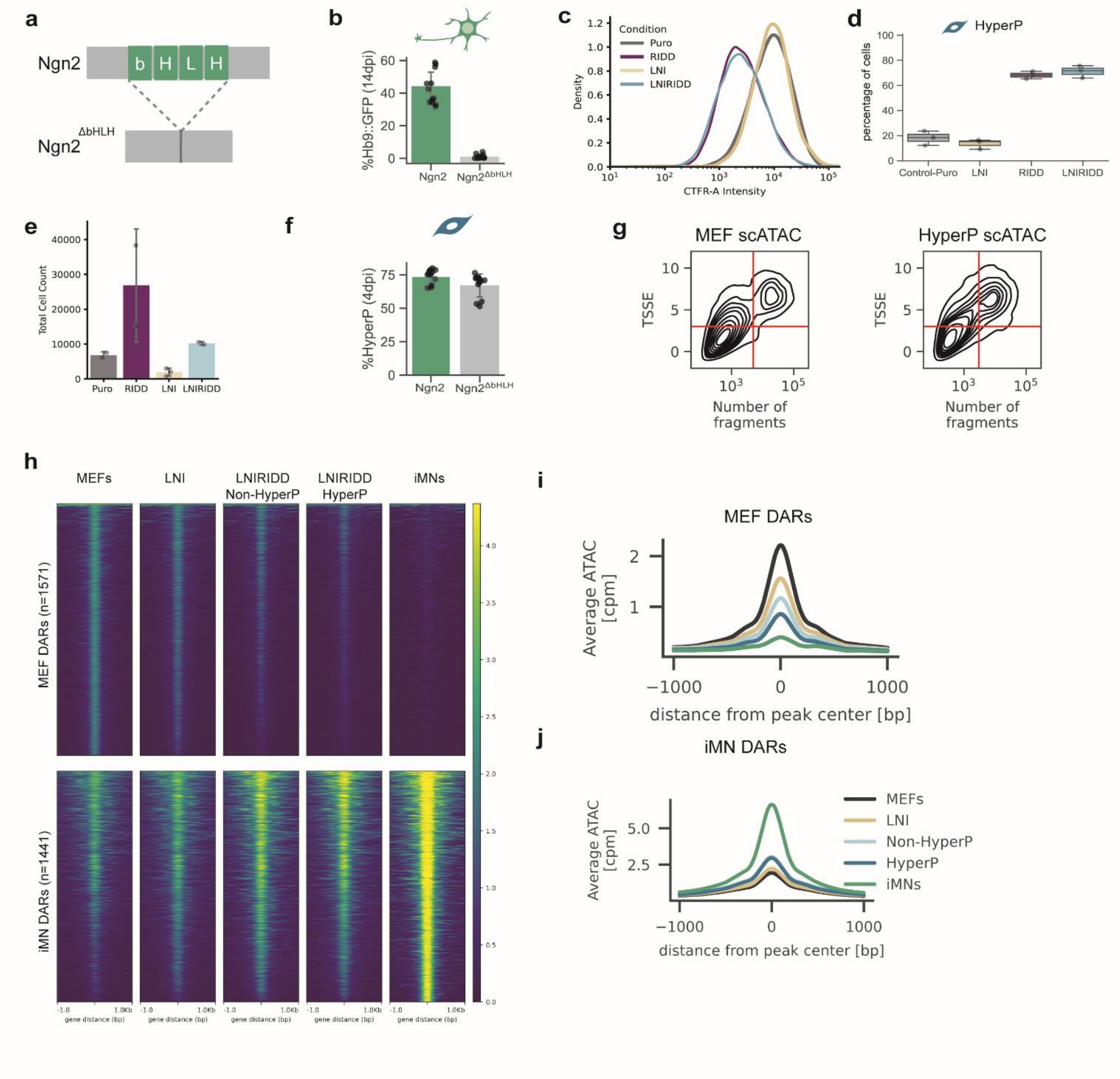


**Fig S1. Reduced accessibility across the axis of conversion defines reprogrammable cells.**

a. Schematic of the bHLH domain in Ngn2 and the mutant bearing the deletion of the bHLH domain.
b. iMN Purity, the fraction of all cells at 14 dpi that are Hb9::GFP positive, at 14 dpi for cells transduced with Lhx3, Isl1, RIDD and the indicated Ngn2.
c. CellTrace Far Red distributions of cells at 4 dpi quantified by flow cytometry.
d. Percentage of cells that are HyperP based on the CellTrace distributions in (c).
e. Quantification of the total number of cells at 4 dpi in each condition via flow cytometry. MEFs were seeded at 10K cells per well of a 96-well plate the day before infection.
f. Percentage of hyperproliferative cells at 4 dpi for cells transduced with Lhx3, Isl1, RIDD and the indicated Ngn2.

g. Kernel density estimate joint plots of transcription start site enrichment (TSSE) and number of unique fragments for single cells in the sci-ATACseq data set for MEFs (top) and LNIRIDD HyperP (bottom). sciATAC datasets were filtered to select the population of cells that clusters in the top right quadrant of these plots.
h. Heatmap showing the read depth normalized chromatin accessibility profiles for each individual DAR in the MEF DAR and iMN DAR sets for all conditions profiled. Data from high quality single cells were merged to create a bulk-accessibility profile.
i. Average read-depth normalized chromatin accessibility signal in MEF DARs for all conditions profiled.
j. Average read-depth normalized chromatin accessibility signal in iMN DARs for all conditions profiled.

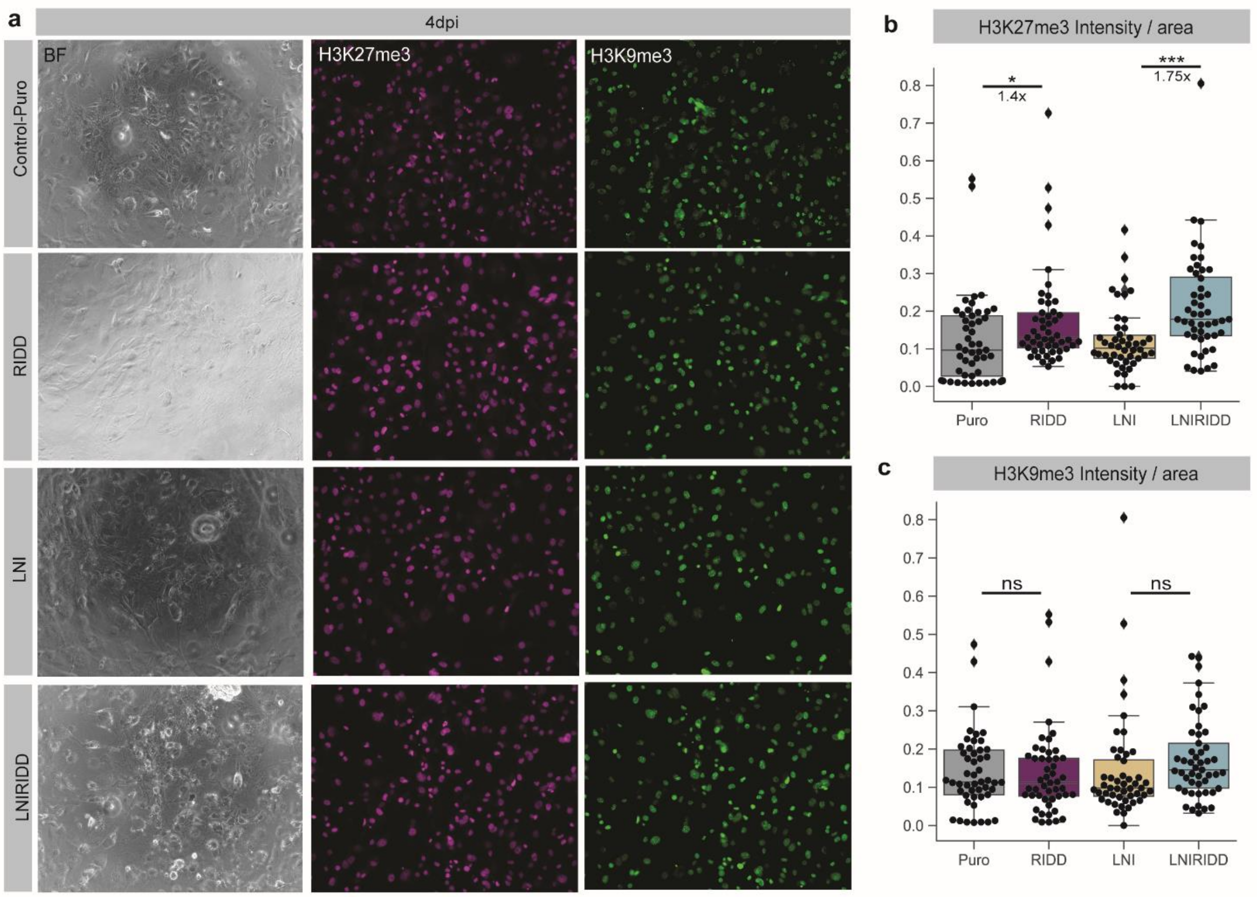


**Fig S2. H3K9me3 levels do not change significantly despite enrichment of H3K27me3 in Hyperproliferative conditions.**

a. Representative brightfield (BF) and immunofluorescence images of cells co-stained for H3K27me3 and H3K9me3.

b-c. Mean fluorescence intensity for H3K27me3 immunofluorescence (b) and H3K9me3 immunofluorescence (c).

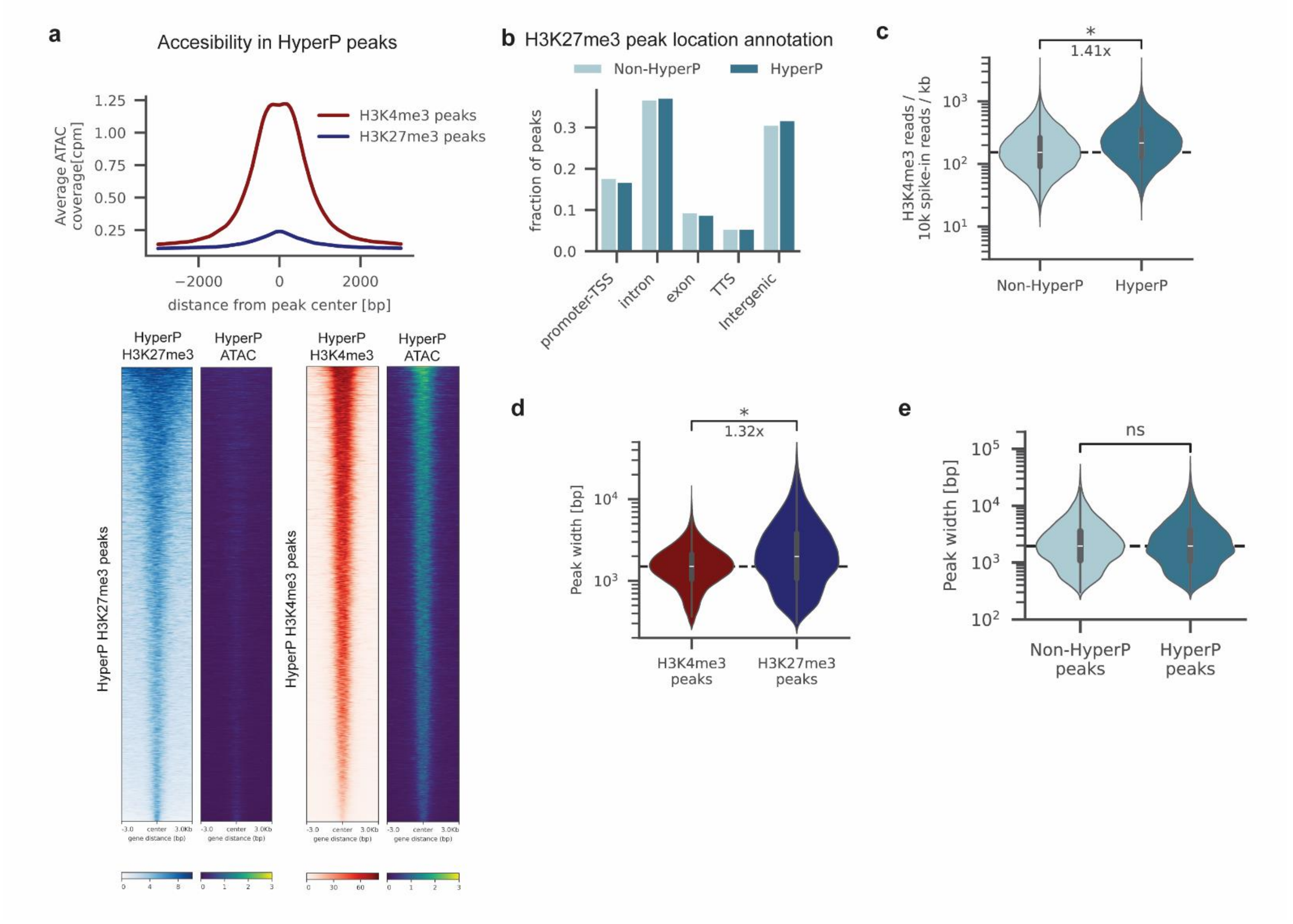


**Fig S3. Hyperproliferative cells broadly enrich H3K27me3 across the genome.**

a. Average accessibility (ATAC counts per million reads [cpm]) in HyperP cells in regions called as H3K27me3 and H3K4me3 peaks in HyperP conditions (top). Heatmaps show CUT&Tag and ATAC normalized read coverage for each region in the two peak sets (bottom).
b. Distribution of peak annotations for annotations for H3K27me3 peaks called in both HyperP and Non-HyperP cells. Annotations were assigned using HOMER.
c. H3K4me3 signal normalized by the number of spike-in reads and region length for peaks called in both HyperP and Non-HyperP conditions. Plot includes data from both biological replicates. Statistical significance determined by one-tailed Kolmogorov–Smirnov test. Fold change indicated is fold change of the median values.
d. Distribution of peak widths in basepairs (bp) for H3K4me3 and H3K27me3 peaks. Statistical significance determined by one-tailed Kolmogorov–Smirnov test. Fold change indicated is fold change of the median values
e. Distribution of peak widths in basepairs (bp) for H3K27me3 peaks in Non-HyperP and HyperP peaks. Lack of statistical significance determined by two-tailed Kolmogorov–Smirnov test.

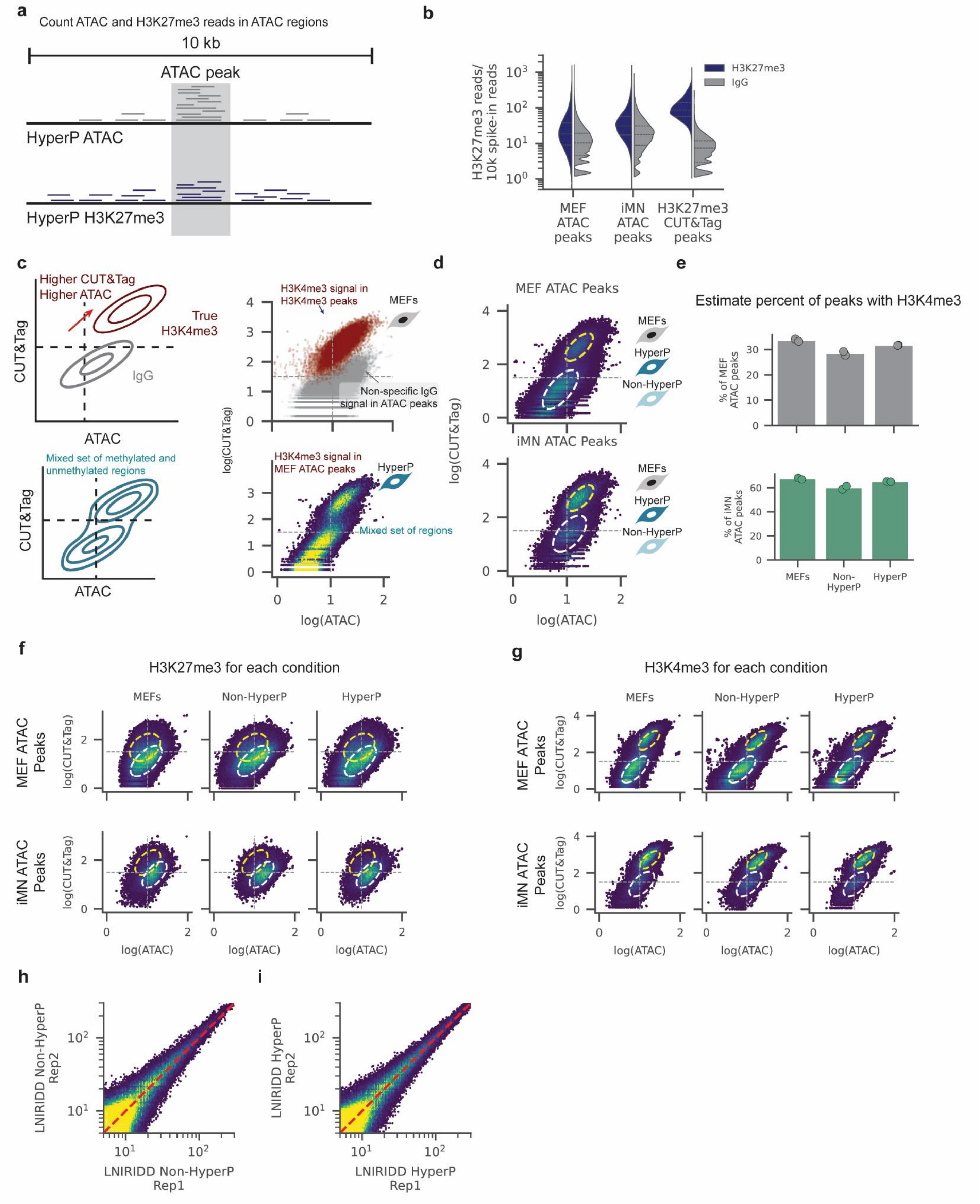
a
Count ATAC and H3K27me3 reads in ATAC regions
10 kb
ATAC peak
HyperP ATAC
HyperP H3K27me3
b
H3K27me3 reads/ 10k spike-in reads
H3K27me3
IgG
MEF ATAC peaks
iMN ATAC peaks
H3K27me3 CUT&Tag peaks
c
Higher CUT&Tag Higher ATAC
True H3K4me3
IgG
CUT&Tag
ATAC
Mixed set of methylated and unmethylated regions
H3K4me3 signal in H3K4me3 peaks
MEFs
Non-specific IgG signal in ATAC peaks
H3K4me3 signal in MEF ATAC peaks
HyperP
Mixed set of regions
log(CUT&Tag)
log(ATAC)
d
MEF ATAC Peaks
iMN ATAC Peaks
MEFs
HyperP
Non-HyperP
e
Estimate percent of peaks with H3K4me3
% of MEF ATAC peaks
% of iMN ATAC peaks
MEFs
Non-HyperP
HyperP
f
H3K27me3 for each condition
g
H3K4me3 for each condition
MEF ATAC Peaks
iMN ATAC Peaks
h
LNIRIDD Non-HyperP Rep2
LNIRIDD Non-HyperP Rep1
i
LNIRIDD HyperP Rep2
LNIRIDD HyperP Rep1

**Fig S4. Gaussian Mixture Models for accounting for high background CUT&TAG signal in accessible regions**

a. As H3K27me3 is a “broad” histone mark, with peaks spanning more than 10 kb, we investigated 10 kb regions centered on regions defined as accessible (ATAC peaks) in MEFs or iMNs. To investigate the relationship between histone marks and accessibility in these regions, we quantified the CUT&Tag signal (number of reads overlapping the regions, normalized by the number of spike-in reads) for each antibody tested and the ATAC signal (number of ATAC transpositions overlapping the region, normalized by the number of ATAC reads).
b. Violin plot showing H3K27me3 CUT&Tag signal and IgG CUT&Tag signal in MEFs for three sets of genomic regions: MEF ATAC peaks, iMN ATAC peaks, and H3K27me3 peaks.
c. Expected correlation for a set of peaks that have H3K4me3 signal above background (top, left). “True” H3K4me3 should have high H3K4me3 signal and high ATAC signal while CUT&Tag signal from an experiment using a non-specific IgG antibody should correlate with accessibility. Scatter plot of and H3K4me3 signal vs accessibility in H3K4me3 peaks in MEFs and IgG CUT&Tag signal vs accessibility in MEFs for regions defined as accessible in MEFs (MEF ATAC peaks) (top, right). When examining a pre-defined set of regions, the set may contain regions with only background CUT&Tag signal and regions with H3K4me3 signal above background (bottom, left). A mixed population is observed in the set of regions surrounding MEF ATAC peaks in HyperP cells (bottom, right)
d. H3K4me3 CUT&Tag signal vs ATAC signal for ATAC peaks after pooling data from MEFs, LNIRIDD HyperP, and LNIRIDD Non-HyperP conditions used to train a Gaussian Mixture Model for the set of 10 kb regions centered on MEF (top) or iMN (bottom) ATAC peaks. The yellow ellipse corresponds to the Gaussian with higher H3K4me3 CUT&Tag signal than background.
e. The fraction of each set of peaks estimated to be in the H3K4me3 high population (yellow ellipses in d) for MEF ATAC peaks (top) or iMN ATAC peaks (bottom). After training the Gaussian Mixture Model on the pooled data, the weight of the two Gaussians were re-optimized for each replicate for each condition to estimate the fraction of peaks within a single CUT&Tag dataset with high H3K4me3 signal. Bar represents the mean of the replicates, with individual replicates plotted as points
f. Distribution of H3K27me3 CUT&Tag signal vs ATAC signal for condition and set of regions. Ellipses represent the two Gaussians in the two-component Gaussian Mixture Model and are consistent within each peak set.
g. Distribution of H3K4me3 CUT&Tag signal vs ATAC signal for condition and set of regions. Ellipses represent the two Gaussians in the two-component Gaussian Mixture Model and are consistent within each peak set.
h. Joint distribution of H3K27me3 CUT&Tag signal in bins for replicates (rep1, rep2) of the LNIRIDD HyperP condition. Signal is quantified as the spike-in normalized read count within a genomic bin. Red dashed line represents the x=y line.
i. Joint distribution of H3K27me3 CUT&Tag signal in bins for replicates (rep1, rep2) of the LNIRIDD Non-HyperP condition. Signal is quantified as the spike-in normalized read count within a genomic bin. Red dashed line represents the x=y line.

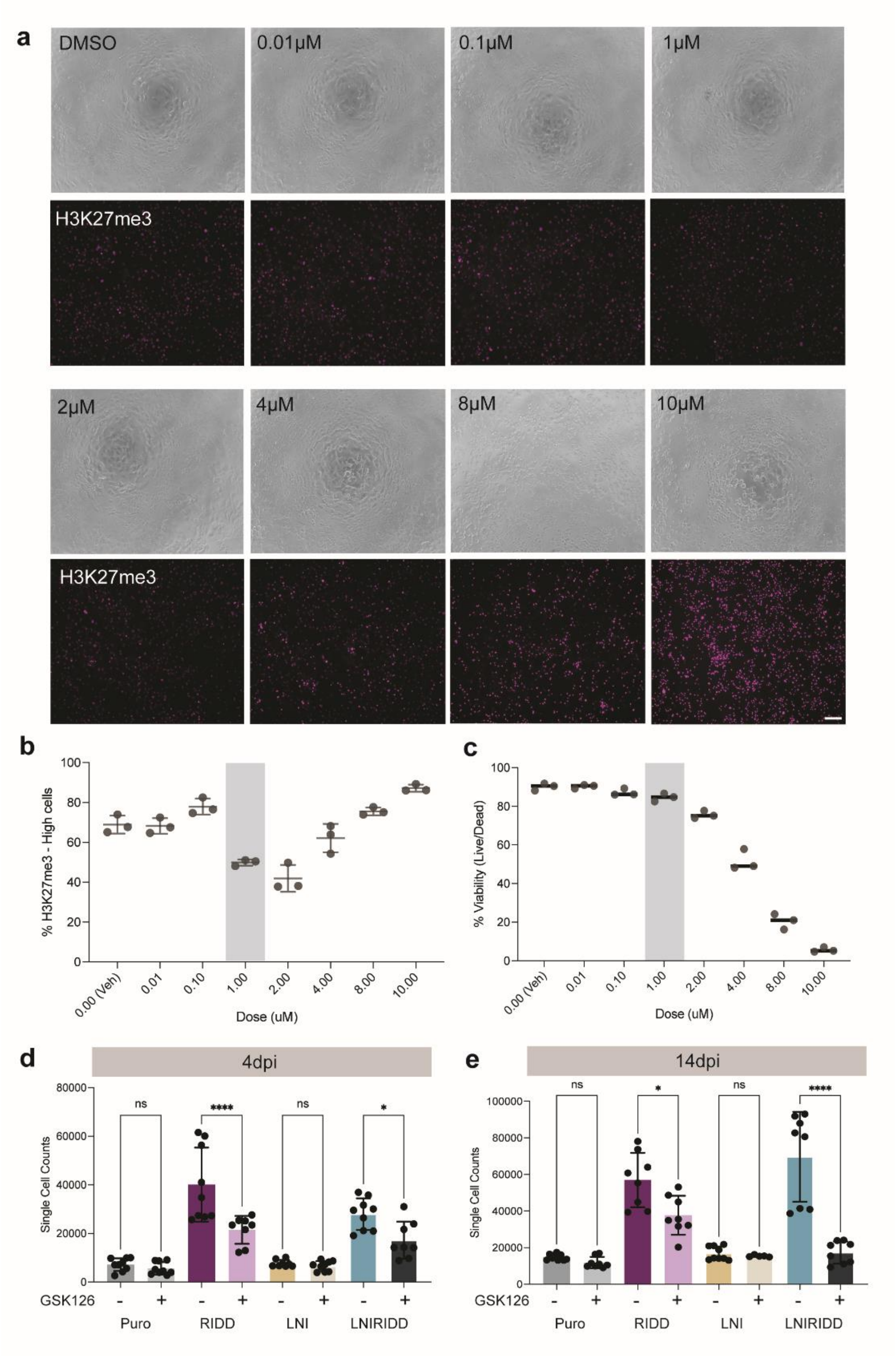


**Fig S5. GSK126 inhibition dose-response and selective effect for Hyperproliferative conditions**

a. Representative brightfield and immunofluorescent images of H3K27me3 staining following treatment with varying concentrations of GSK126. Treatment above 2 μM for 24 hours in MEFs induces autofluorescence and cell detachment.
b. Fraction of MEFs with high levels of H3K27me3 when treated with the indicated concentration of GSK126 for 24 hours. Quantified by flow cytometry. Shaded region indicates chosen dose for conversion experiments.

c. Fraction of MEFs that are viable when treated with the indicated dose of GSK126 for 24 hours. Cells were stained with the Zombie NIR Live/Dead stain. Dead cells are stained by the Zombie NIR Live/Dead stain, viable cells are negative for the stain. Shaded region indicates chosen dose for conversion experiments.
d. Quantification of the total number of cells at 4 dpi in each condition via flow cytometry, with and without acute treatment with GSK126 from 1-2 dpi. MEFs were seeded at 10K cells per well of a 96-well plate the day before infection.
e. Quantification of the total number of cells at 14 dpi in each condition via flow cytometry, with and without acute treatment with GSK126 from 1-2 dpi. MEFs were seeded at 10K cells per well of a 96-well plate the day before infection.

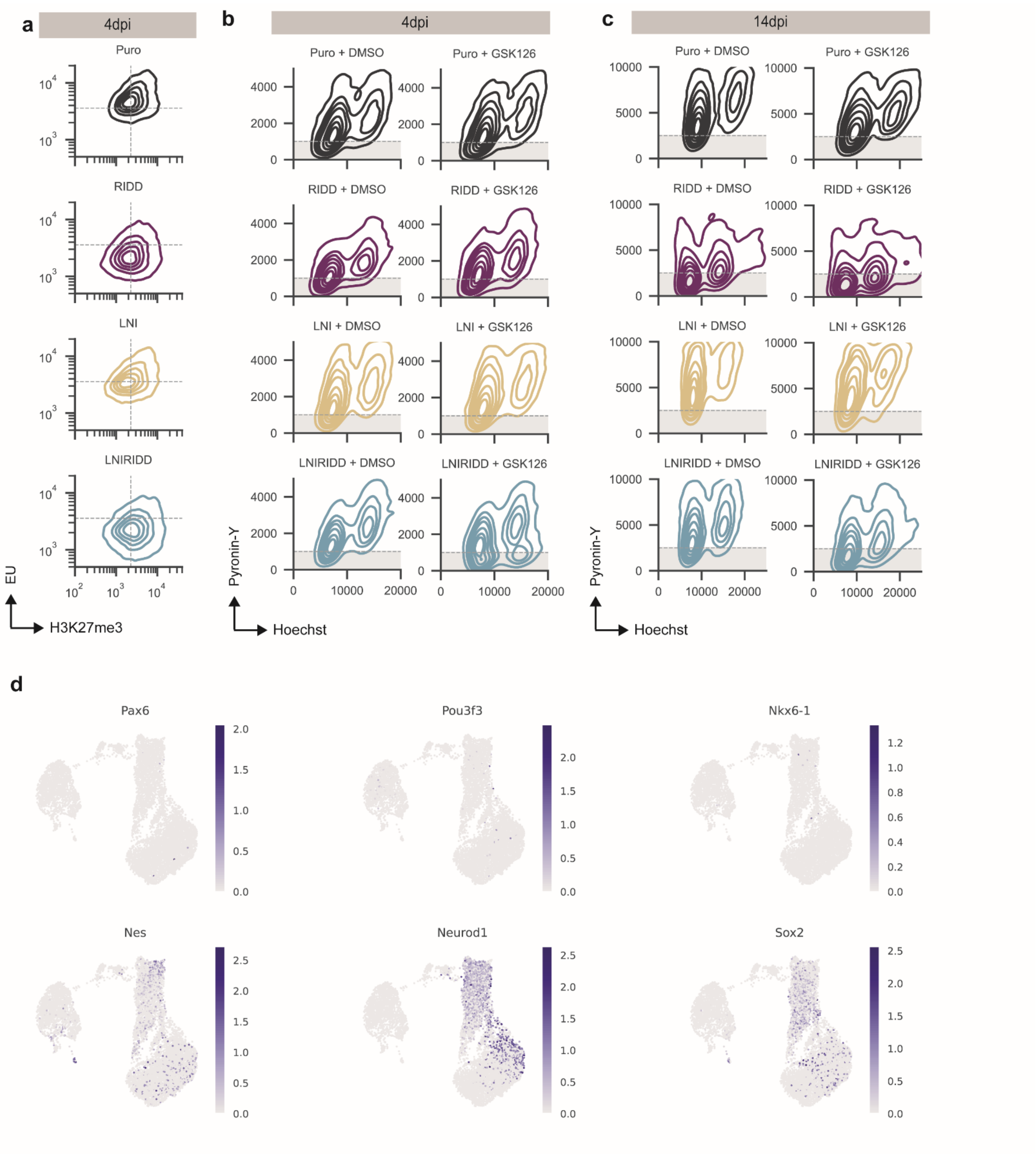


**Fig S6. Low RNA and Reduced relative transcription, LNIRIDD HyperP cells are not enriched for neural progenitor markers**

a. Kernel density estimate joint plot of cells stained for H3K27me3 and EU for the indicated condition quantified by flow cytometry at 4 dpi.

b. Kernel density estimate joint plot of cells stained for Pyronin-Y and Hoechst for the indicated condition with acute DMSO or GSK126 treatment from 1-2dpi, quantified by flow cytometry at 4 dpi.
c. Kernel density estimate joint plot of cells stained for Pyronin-Y and Hoechst for the indicated condition with acute DMSO or GSK126 treatment from 1-2dpi, quantified by flow cytometry at 14 dpi.
d. Log expression of the indicated progenitor-cell associated gene projected onto the UMAP of the scRNAseq data.

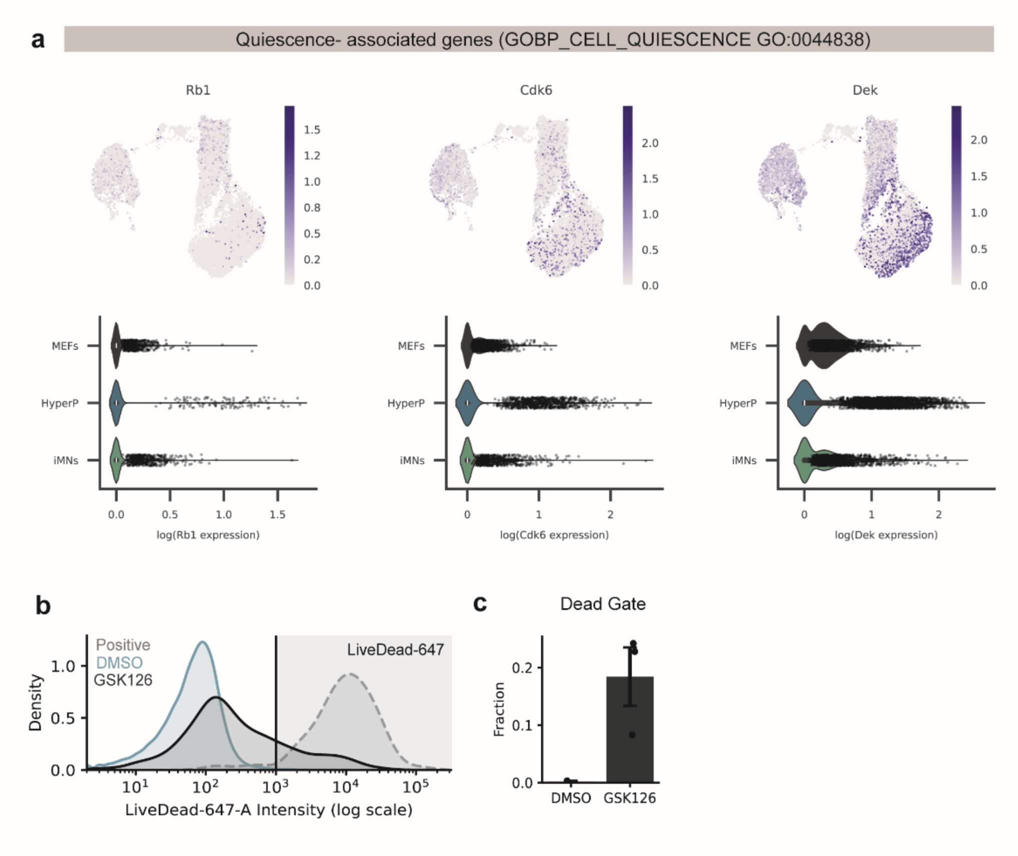


**Fig S7. Cells transit via a quiescent-like state protected by H3K27me3.**

a. Log expression of the indicated quiescence-associated gene projected onto the UMAP of the scRNA-seq data (top) and distribution of single cell gene expression (bottom). Dots on violin plots represent expression for any cell with expression greater than 0 for the indicated gene.
b. Kernel density estimate plot of the single cell distribution of Zombie-NIR Live/Dead stain fluorescence for LNIRIDD cells at 4 dpi. Dead cells are those stain positive for the Live/Dead stain (grey region).
c. Fraction of cells that stain positive for the Zombie-NIR Live/Dead stain in LNIRIDD cells at 4 dpi.
